# A Simulation Study on High Spatio-Temporal Resolution Acousto-Electrophysiological Neuroimaging

**DOI:** 10.1101/2023.06.07.544012

**Authors:** Ruben Schoeters, Thomas Tarnaud, Luc Martens, Emmeric Tanghe

## Abstract

Acousto-electrophysiological neuroimaging is a technique hypothesized to record electrophysiological activity of the brain with millimeter spatial and sub-millisecond temporal resolution. This improvement is obtained by tagging areas with focused ultrasound (fUS). Due to mechanical vibration with respect to the measuring electrodes, the electrical activity of the marked region will be modulated onto the ultrasonic frequency. The region’s electrical activity can subsequently be retrieved via demodulation of the measured signal. In this study, the feasibility of this hypothesized technique is tested. This is done by calculating the forward electroencephalography (EEG) response under quasi-static assumptions. The head is simplified as a set of concentric spheres. Two sizes are evaluated representing human and mouse brains. Moreover, feasibility is assessed for wet and dry transcranial, and for cortically placed electrodes. The activity sources are modeled by dipoles, with their current intensity profile drawn from a power-law power spectral density. It is shown that mechanical vibration modulates the endogenous activity onto the ultrasonic frequency. The signal strength depends non-linearly on the alignment between dipole orientation, vibration direction and recording point. The strongest signal is measured when these three dependencies are perfectly aligned. The signal strengths are in the pV-range for a dipole moment of 5 nAm and ultrasonic pressures within FDA-limits. The endogenous activity can then be accurately reconstructed via demodulation. Two interference types are investigated: vibrational and static. Depending on the vibrational interference, it is shown that millimeter resolution signal detection is possible also for deep brain regions. Subsequently, successful demodulation depends on the static interference, that at MHz-range has to be sub-picovolt.

## I. Introduction

Functional neuroimaging techniques are well established in the medical world. They are used in disease diagnosis, monitoring and for signal acquisition in brain computer interfaces [1]–[3]. The different methods can be categorized into two groups based on the measured activity: direct electrophysiological and indirect hemodynamic [4].

Functional magnetic resonance imaging (fMRI), functional near-infrared spectroscopy (fNIRS) and positron emission tomography (PET) are modalities among the latter group. They are based on the neurovascular coupling, where changes in neuronal activity result in, for instance, altered blood flow or metabolic changes [3]. Compared to the electrophysiological measuring modalities, they have typically a better spatial resolution but a worse temporal resolution. fMRI performs best on spatial resolution. This is in the mm-range for reasonable acquisition time (1-2 s) and full brain coverage. It can even go sub-mm, however, at the cost of temporal resolution, tissue coverage or the need for complex infrastructure [5], [6]. PET has a slightly lower spatial resolution and a temporal resolution up to minutes. An important advantage of both techniques is their ability to map activity of deep regions. fNIRS is more restricted to the cortex with a maximal depth of 1-3 cm below the skull. The spatial resolution is also lower with respect to PET and fMRI, being in the upper mm-range (5-10 mm). On the other hand, the sampling rates are around 10 Hz [1], [4], [7], [8].

Electroencephalography (EEG) and Magnetoencephalography (MEG) directly and non-invasively measure the electrophysiological activity. Their spatial resolution is in general lower than that of the hemodynamic modalities. Due to the differences in electrical properties between biological tissues, such as scalp and skull, the EEG-signal strongly smears out onto a larger scalp area (*>*10 cm^2^ [9], 3 cm^2^ [10]) [11]. The differences in magnetic permeability of the tissues are negligible resulting in less dispersion. Consequently, it is argued that MEG performs better in source localization with 1 cm and 2.5 cm error for MEG and EEG, respectively [12]. On the other hand, the study of Klamer et al. (2015) [13] showed a significantly lower localization error for high-density EEG (256 channels) compared to MEG (275 channels) when individual head models were used. Thus, by using realistic models to overcome the dispersion, better localization can be obtained with EEG than with MEG for the same number of sensors [13], [14]. The electropyhsiological activity can also be measured with implanted electrode arrays. This can provide local information of a small volume of cells. Of course, they are invasive, causing tissue damage. Furthermore, electrochemical reactions can affect signal stability [4]. Finally, a less invasive approach is electrocorticography (ECoG) where electrodes are subdurally implanted. By surpassing the skull, the spatial resolution is improved with respect to EEG (5 mm) [9]. However, the coverage is restricted to the region below the electrodes [15], [16]. All these electrophysiological modalities have a superior sub-millisecond temporal resolution [12]. Furthermore, they directly measure neuronal acitivity, while there is a latency of ∼ 5 s between neuronal activity and hemodynamic changes [1], [7]. Moreover, a medio-lateral shift between hemodynamic and electrophysiological localization has been observed, as well [13].

A single technique’s disadvantage could be overcome by adopting a multimodal approach. For instance, fMRI and EEG have strong complementary strengths. A hurdle, however, is the concurrent electromagnetic interference inducing artifacts [3]. On this aspect fNIRS is promising because it can be coupled with EEG, MEG or fMRI seeing it does not make use of metallic probes [7]. Although often thought as competing, also EEG and MEG provide complementary information allowing for better accuracies [11], [12].

Other important properties are cost, equipment size, and patient comfort, the latter including, portability, noise and movement freedom. Here, EEG scores the best (as well as fNIRS), making it together with its high temporal resolution an attractive and therefore highly investigated technique [4], [8]. Advances in electrophysiological source imaging (ESI) have significantly improved the localization errors of EEG. The goal is to find the underlying sources responsible for the measured EEG signal, by solving the inverse problem. This problem is strongly ill-posed due to the in theory billions of sources and only limited number of electrodes. A summary of all contributing sources to the electric field is given in [9]. A solution to this inverse problem is found by minimizing the difference between measured and calculated signals, the latter obtained by solving the forward problem. A review on the different ESI-algorithms is given in He et al. (2018) [3] and Asadzadeh et al. (2020) [17]. It has to be noted that centimeter differences in localization depending on the used ESI method exist [18], [19]. The accuracy can be improved by increasing the number of electrodes. However, the additional absolute improvement decreases with increasing number of electrodes. Also, the localization error increases with depth, while the rate drops with a higher number of electrodes [20].

The chosen forward head model strongly affects the localization accuracy, as well. It translates the activity of a source in the brain to the electrode. As aforementioned, the signals are strongly dispersed by differences in electrical properties between biological tissues. Realistic head models can improve accuracy by including anisotropy and inhomogeneity of the tissues [17], [18], [21]. Klamer et al. (2015) [13] also demonstrated a significant inter-subject variability. Consequently, a patient specific head model needs to be constructed, which can be obtained via magnetic resonance image of the head. On the other hand, this increases computational complexity preventing real-time analysis. Also the need for an MRI scan impacts the ease-of-use of EEG.

Aside from the conventional techniques discussed above, the use of ultrasound for functional neuroimaging is receiving increased attention. Functional ultrasound imaging is another hemodynamic technique with high spatial and temporal resolution (millisecond and millimeter). Although limited to two-dimensional imaging planes, transition to 4D image acquisition is being investigated [22]. Alternatively, by probing the brain with focused ultrasound (see figure 1 (A)), it has been hypothesized that the electrophysiological activity in the ultrasound focus will be modulated onto the ultrasound frequency. The endogenous signal can be retrieved after demodulation. In this manner a superior spatio-temporal resolution can be obtained, where the spatial resolution depends on the size of the ultrasound focal zone (mm-range) and the temporal resolution is sub-milisecond (like with EEG). Moreover, no inverse problem needs to be solved, removing the hurdles mentioned above. Bin He (2016) [23], postulated that modulation is achieved by relative motion of the source with respect to receiver due to mechanical vibration induced by the ultrasonic field. This was tentatively called acousto-electrophyiological neuroimaging (AENI). Another possible underlying mechanism is the acousto-electric effect (AE), i.e., a pressure induced change in conductivity [24].

**Fig. 1.**
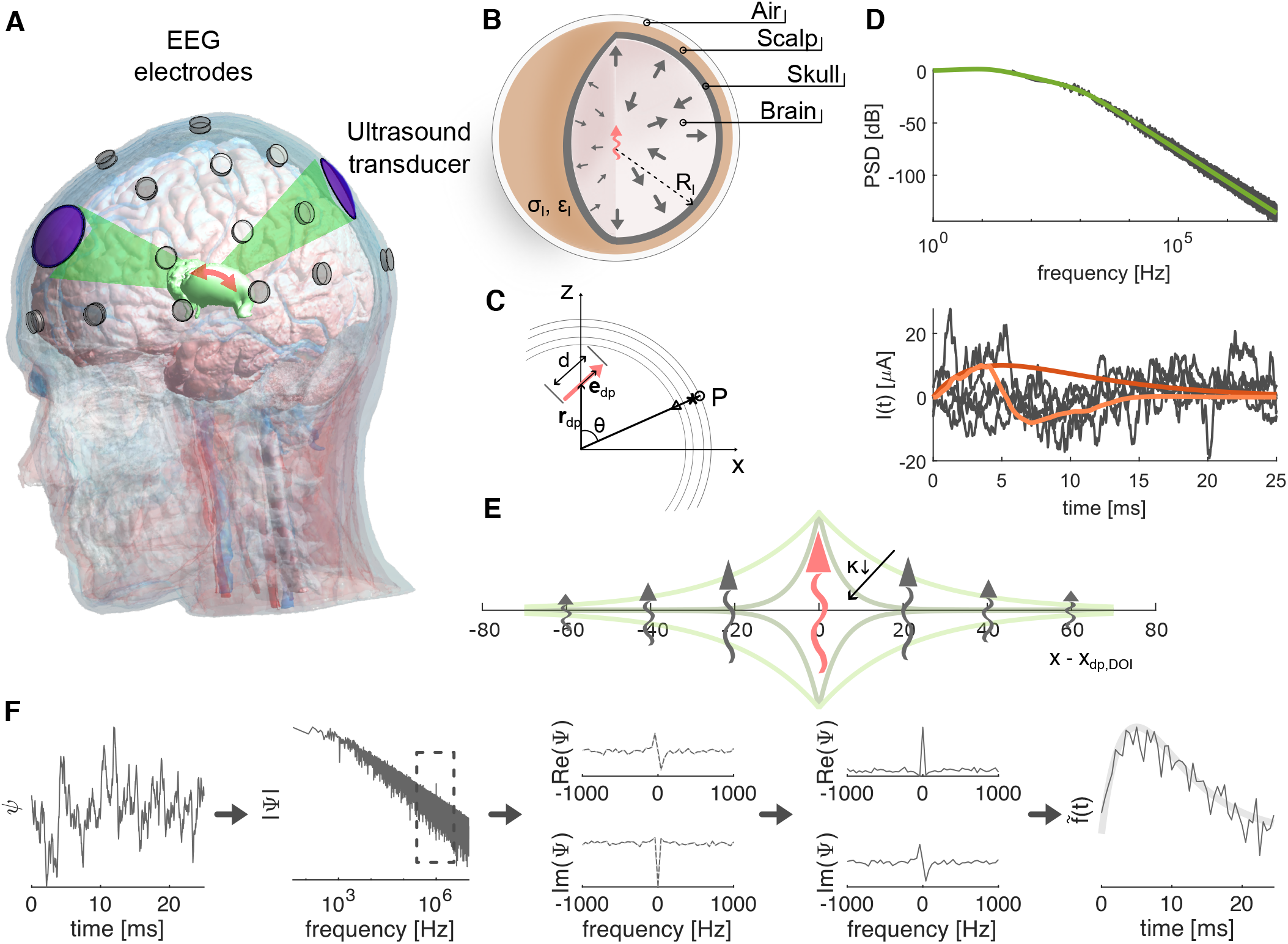
Concept of acousto-electrophysiological neuroimaging and description of used methods. **A** A highly focalized ultrasonic field is applied to the brain to mark a specific region. Mechanical vibration of the tagged region modulates the biological signal on the ultrasound’s carrier frequency, spatially encoding the electrical brain activity. Demodulation of the measured EEG signal should return the marked region’s activity. **B** The spherical four-layer head model, with each layer specified by its conductivity *σ*_*l*_, permittivity *ϵ*_*l*_ and radius R_*l*_. The electrical activity of different brain regions are represented as dipoles (arrows) with the tagged region in red. **C** Schematic of the equivalent dipole and its parameters. The potential is measured at P in the xz-plane. The circle, asterisk and triangle indicate the corresponding positions of the dry, wet and cortical electrodes. **D** The time dependent current intensities of the different dipoles. Top, the power spectral density (PSD) profile from which the time signals are reconstructed (green line without noise). Below, the time dependent signals, with red and orange (i.e., respectively, alpha-function and alpha-train) the time signal of the tagged dipole and the gray lines the signal of the background dipoles derived from the PSD-profile above. **E** The one dimensional applied displacement field for increasing values of the spatial constant *κ*. **F** The signal demodulation protocol. From left to right: Fourier transform, bandpass filter and demodulation to baseband, zero order detrend followed by phase shift correction and finally inverse Fourier transform.

The latter group (Witte et al. [25]) already performed numerous in-vitro experiments, proving that the electric signal gets modulated onto the ultrasound carrier [26]. They showed that this technique could be used for 4D ultrasound current source density imaging (UCSDI) modality [27], [28]. The spatial resolution was indeed in line with the full width at half maximum of the US-focal zone. When applied to the brain this was termed transcranial acoustoelectric brain imaging (tABI) [29], [30]. They confirmed that a broadband EEG-like signal from deep within the brain (40 mm) could be retrieved [30]. In this simulation study, the mechanism as stated by He (2016) [23] is investigated. The head is modeled as a set of concentric spheres. Current dipoles are used to represent the electrophysiological activity generated by a small volume of neurons. Subsequently, the EEG response is determined by solving the general Poisson equation in spherical coordinates under quasi-static assumptions. The feasibility is tested for two head models (i.e., human and mouse) and three electrode types (i.e., scalp wet and dry, and cortical electrodes). It is shown that relative motion induced by mechanical vibration does modulate the endogenous signal onto the used ultrasonic frequency. The maximal signal strengths are determined together with the optimal dipole orientation, displacement direction and electrode position. Furthermore, two interference categories are identified. The vibrational interference originates from neighboring vibrating regions, accountable for the achievable spatial resolution. And, the static interference, that is the endogenous biological signal noise at high frequencies, responsible for the technique’s feasibility. Finally, the necessary level of dipole moment power of a single dipole’s current intensity profile in order for the technique to be safe is determined.

## II. Methods and Materials

The framework used to test the hypothesized method is described below. A quasi-static approach is adopted to model the displacement of the brain regions. We opted for a simplified solution of the problem for mathematical convenience, i.e., the head is modeled as a set of concentric spheres. Next, we elaborate on the dipole moments and their time dependent signals, important for the static interference contribution. A synthetic ultrasonic field is applied to control the influence of vibrational interference. Both are biological noise sources that interfere with the signal of interest. The former originates from the static field. The latter is due to vibrating regions not being the region of interest. Finally, the signal demodulation method, metrics and FDA limits, used to test the method’s feasibility are described. The full study was performed in MATLAB R2021b.

### A. Quasi-Static Electromagnetic Field

Application of the ultrasonic field causes brain regions to move with respect to the EEG-electrodes. The conventional static solution for the forward EEG problem will not suffice. A quasi-static electromagnetic field approximation is selected. The decision is based on an order of magnitude analysis using the Liénard-Wiechert fields (see supplementary section II). The analysis showed that quasi-static electric potential contributions dominate all relativistic components (i.e., due to the Doppler effect, source acceleration and finite propagation speed of light) for an angular ultrasonic frequency:

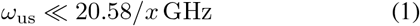

with *x* the distance between observer and oscillating source in cm. Under quasi-static assumptions, the induced electric potential (*Ψ*) for a displacement in the x-direction is:

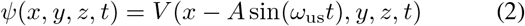

with A the displacement amplitude, *ω*_us_ the angular ultrasonic frequency and *V* the induced electric potential in the reference frame of the source. A Taylor series for A ≪ 1 yields:

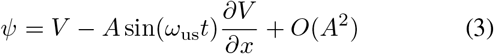

Consequently, the measured signal contains the information of the source near the applied ultrasonic frequency. In other words, it is modulated onto *f*_us_.

### B. Spherical Four-Layer Head Model

The head is modeled as a set of concentric homogeneous spheres, figure 1 (B). An analytical solution can be obtained by solving the general Poisson equation in spherical coordinates. For a current dipole with moment *I*_max_ *f*(*t*)**d**_dp_ on the z-axis, the induced electric potential *V*_*l*_ in volume *l* for an arbitrary point on the XZ-plane can be determined via:

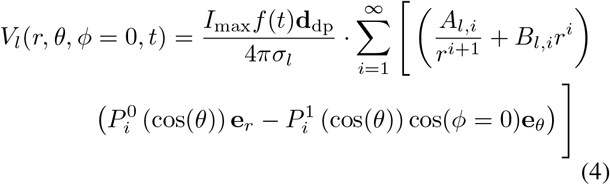

where *σ*_*l*_ is the conductivity of volume *l. r, θ* and *ϕ* are spherical coordinates. The terms 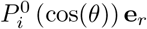 and 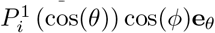 correspond to radial and tangential components of the dipole, respectively, with 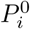 and 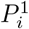 the associated Legendre polynomial of zeroth and first order [31], [32]. The Condon-Shortley phase is included in calculating the associated Legendre polynomials, explaining the minus sign between radial and tangential dipole terms (unlike in Arthur and Geselowitz (1970) [31]).

In this study, three electrode types are of interest, i.e., cortical, and wet and dry scalp electrodes. Although, no actual electrodes are modeled, these types can be associated with different positions in the head. Therefore, the model consists of four concentric shells, each defined by an outer radius *R*_*l*_. From inside to outside: the first is brain matter, followed by the skull, next the scalp and finally a layer of air. For the cortical electrode, the point of interest is at the brain skull boundary (*R*_brain_). The wet scalp measurement occurs at the scalp air boundary (*R*_scalp_). The dry scalp is inside the air shell (*R*_air_). Under the following conditions, a solution for eq. (S.3) can be found.

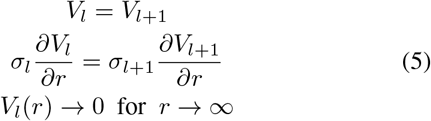

Due to the interest in multiple radial positions, the solution to eq. (S.3) cannot be simplified. Therefore, it is not depicted here but can be found in supplementary information, section III. The radius of each shell can be found in table I [21]. The number of terms included in the numerical simulation in eq. (S.3) is regulated by an absolute and relative tolerance, i.e., 10^−13^ and 10^−10^, respectively, limited by a maximum of 5000 terms.

**Table I.**
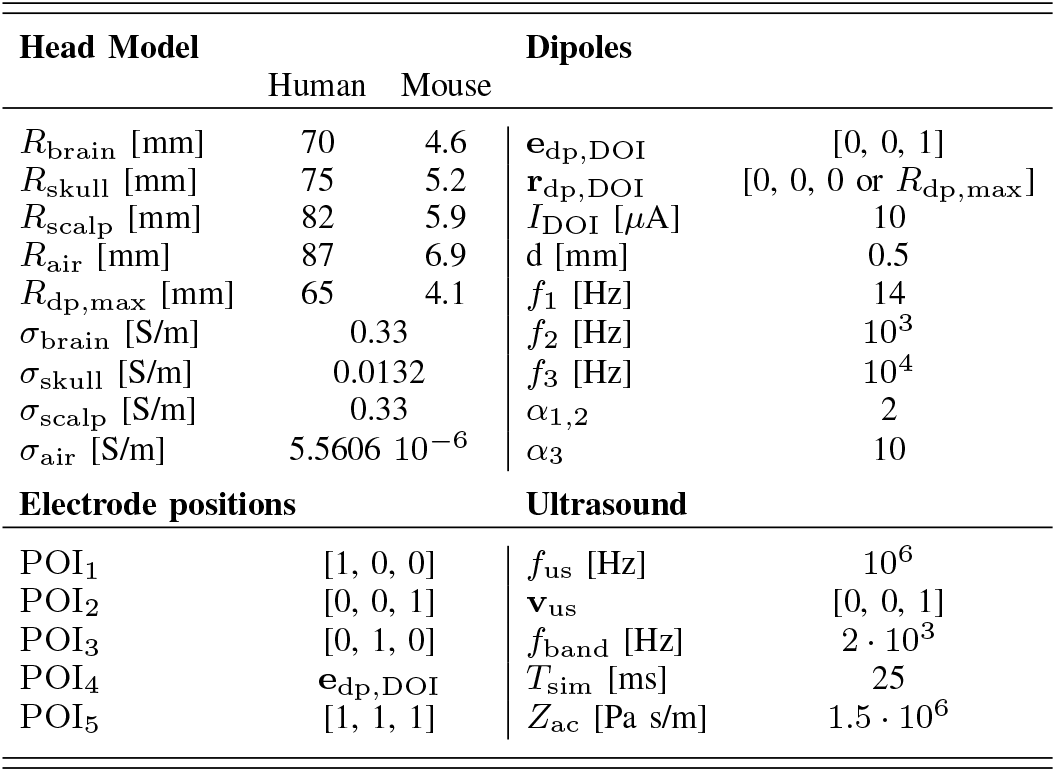
Summary of all parameters values. Unless otherwise specified, these values are used in each simulation.

Due to the interest of the signal at *f*_us_ *± f*_band_*/*2, the used conductivities are the modulus of the complex conductivities 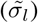 near *f*_us_. At *f*_us_, the capacitive effects are negligible for the brain, skull and scalp conductivities. Therefore, 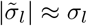. In the air layer, on the other hand, capacitive effects dominate, resulting in 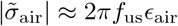 with *ϵ*_air_ the permittivity of air. The values are summarized in table I, taken from the IT’IS database [33].

### C. The Dipole Moment

EEG response is caused by extracellular neuron currents in response to transmembrane currents, also known as secondary return, or volume currents. These currents can be modeled by a multipole expansion [2], [21]. However, typically only the first order expansion is used, i.e., the current dipole. Here, it represents the activity of a small volume of parallel neurons [2], [21], [31], [32]. The dipole is characterized by: its position (**r**_dp_ = [*x, y, z*], typically halfway between the current source and sink), its orientation (defined by unit vector **e**_dp_), the current intensity (*I*) and distance between the monopoles (*d*), see figure 1 (C). The dipole moment density is then

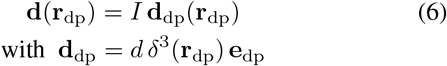

with *d* the Dirac delta. Unless otherwise specified, the dipoles are radially oriented. Moreover, they are uniformly distributed on *n* (= 3, 5 or 10) concentric spheres in the brain. The layers are equidistant, with the most outer layer at *R*_dp,max_, and have the same dipole densities.

The current intensity varies over time. An appropriate time dependent signal needs to be chosen to obtain an estimate of the noise level at *f*_us_. The time dependent signals of any dipole not being the dipole of interest is derived from a power spectrum density profile:

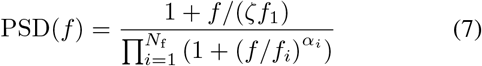

with *f*_*i*_ the *i*th cut-off frequency, *α*_*i*_ the attenuation coefficient and *N*_f_ = 2 or 3, for the vibrational and static interference analysis, respectively. *ζ* is randomly chosen between [0.3, 1]. Consequently, the PSD reaches a maximum between *ζf*_1_ and *f*_1_. Extra noise is added to increase the variance amongst the dipoles. This is drawn from log 𝒩 (0, 0.5). The time varying current intensity is then obtained via:

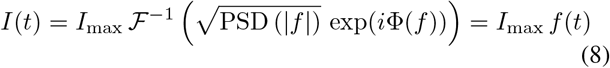

with Φ(*f*) an odd function of random phases, *i* the imaginary unit and ℱ^−1^ the inverse Fourier transform. The maximum current intensities are drawn from 𝒩 (*I*_DOI_, 3) uA. The parameter values are summarized in table I. The time signal for the dipole of interest (DOI, see section II-D) follows an alpha-function eq. 10 or if explicitly specified an alpha-train eq. 11.

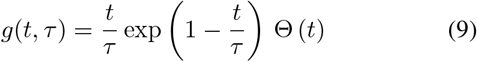

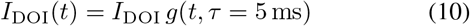

or

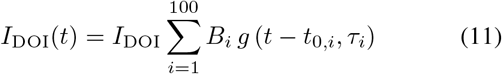

with *τ*_*i*_ drawn from 𝒩 (2, 0.2) ms, *t*_0,*i*_ from |𝒩 (0, 6.5)| ms and *B*_*i*_ from 𝒩 (0, 1.5). Θ(*t*) is the Heaviside step function.

Furthermore, *B*_*i*_ is normalized with respect to the largest absolute *B*_*i*_ value. The maximum dipole moment of the DOI is set to 5 nAm, after the synchronous activity of 10^4^ neurons with a single dipole moment of 0.5 pAm. With a *d* = 500 *μ*m, this results in a *I*_DOI_ = 10 *μ*A [34], [35] The sampling frequency used is 20 times *f*_us_. An example of the applied current intensities is shown in figure 1 (D).

### D. Ultrasonic Field

An synthetic ultrasonic field is used to provide more control of the displacement field distribution.

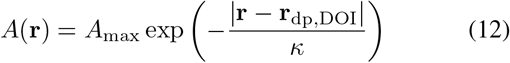

with *κ* the spatial constant in mm. In this study the displacement vector is always in the z-direction. The field and effect of *κ* is illustrated in figure 1 (E). The dipole of interest (DOI) is the dipole at the location where the displacement is maximal (*A*_max_). This is the tagged region.

### E. Signal Processing

The electric potential is measured at 5 locations or positions of interest (POIs). The POIs are defined as the location vector connecting the origin and the measurement point (see table I). Two additional POIs are included as well: mPOI and mPSO. The former being the mean of all POIs, the latter is the mean of POIs that received DOI’s signal within the same order of magnitude of the highest received signal. Depending on the electrode type of interest (see section II-B), the POI is then located at *R*_*l*_ *·* POI_*i*_*/* |POI_*i*_|. Subsequently, the signal at *f*_us_ *± f*_band_*/*2 is demodulated following (figure 1 (F)):

1. Fourier transform: *Ψ* (*POI*_*i*_, *t*) →Ψ (*POI*_*i*_, *f*)
2. bandpass filter (*f*_us_ *± f*_band_*/*2) followed by demodulation to baseband
3. zero order detrend, i.e., subtracting mean from real and imaginary parts.
4. compensate for phase shift due to sine (see eq. (3))
5. inverse Fourier transform

Normalization of the obtained signal results in:

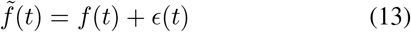

where *f*(*t*) is the normalized time varying current intensity (see eq. (8)) and *ϵ*(*t*) the error due to interference of other dipoles. Finally, the root mean square error (RMSE), with respect to the input signal is determined, to asses the reconstruction accuracy.

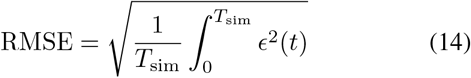

A titration process is used to determine the thresholds of the displacement amplitude *A*_max_ and spatial constant *κ* to obtain a given RMSE-value. This process adopts the bisection method where the midpoint is the logarithmic mean of the interval [*a, b*] (i.e., log^10^ *c* = (log^10^ *a* +log^10^ *b*)*/*2). At the start *b/a* = 10. To limit computation time only a max of 5 iterations are evaluated.

### F. FDA Limits

The FDA limits are used to asses the methods feasibility [36].

- Pulsed average intensity:

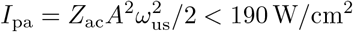
- Temporal average intensity:

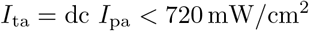
- Mechanical index:

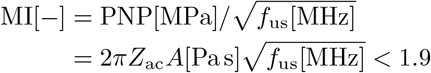

with *Z*_ac_ the specific acoustic impedance, PNP the peak negative pressure in [MPa] and dc the duty cycle. Here, a plane wave approximation is applied in the focus (*p*_us_*/v*_us_ = *Z*_ac_), which is valid under the assumption of a small transducer convergence angle [37].

## III. Results

Below the results of this simulation study are shown. First the measurable signal strength of a single dipole is investigated. This led to a preferred displacement direction and dipole orientation combination that is further maintained in following subsections. Next, two different interference sources are addressed. The vibrational interference is investigated followed by the static interference. Finally, the importance of the imposed reconstruction accuracy and dipole current waveform is analyzed.

### A. Measurable Signal Strength of Single Vibrating Dipole

The relationship between dipole position (**r**_dp_), orientation (**e**_dp_), measurement position (POI) and displacement direction (**e**_z_) is investigated first. A single dipole is used, with a constant dipole moment of 5 nAm. The dipole is moved radially (*r*_dp_) from 0 to *R*_dp,max_. Moreover, it is rotated in the yz-plane for a varying polar angle (*θ*_dp_) from 0 to *π* radians. At each position, the dipole is oriented radially outwards (**e**_dp_ = **e**_r_). The displacement amplitude (*A*_max_) is 10 nm. The results are shown in figure 2.

**Fig. 2.**
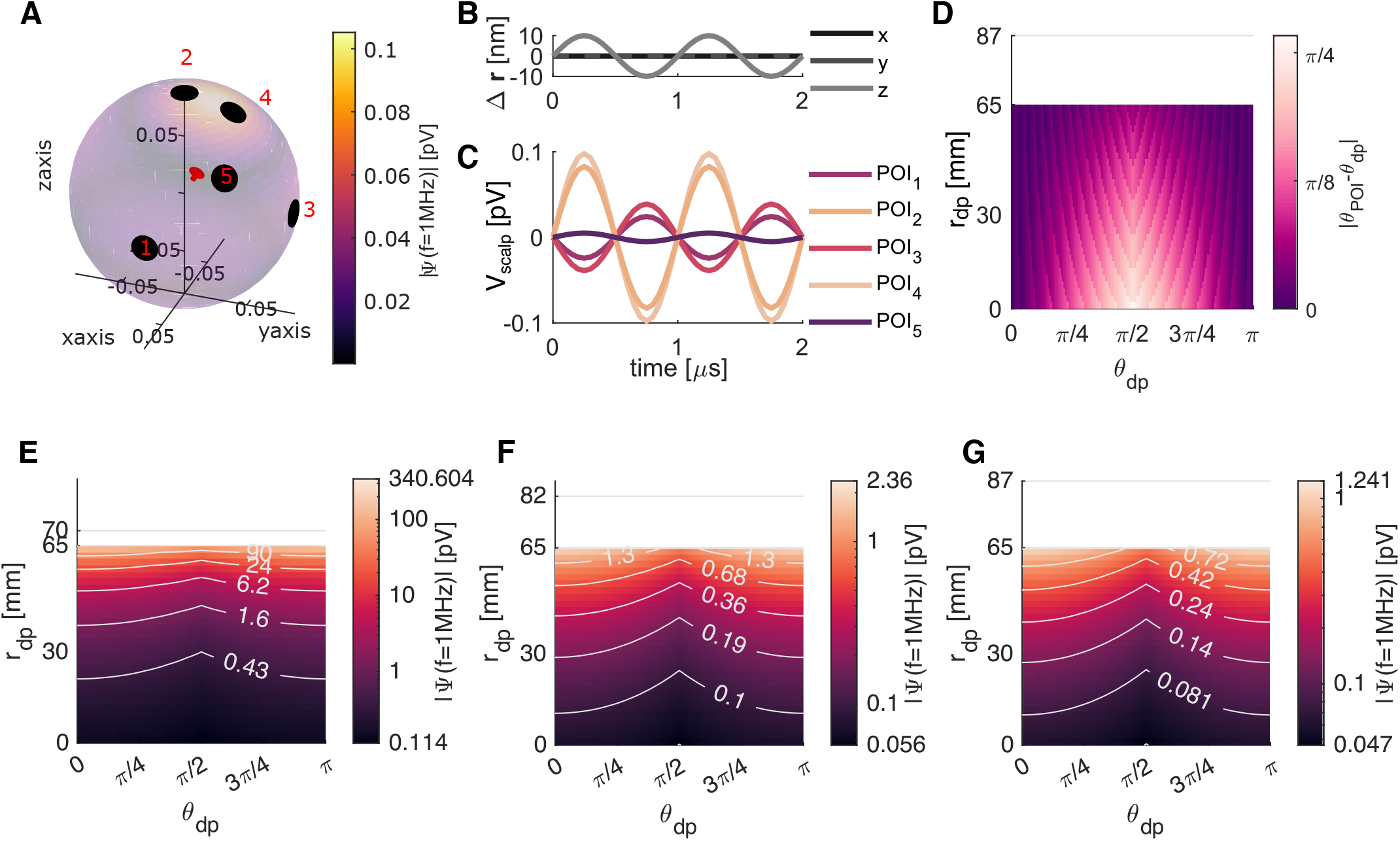
Measurable signal strength for single dipole with constant dipole moment of 5 nAm. The dipole is located in the yz-plane, and is moved radially (*r*_dp_) and with changing co-elevation (*θ*_dp_). **A** Distribution of the amplitude of the *f*_us_ component measurable at the scalp-air interface (*R*_scalp_). The tagged dipole is indicated by a red arrow. The point of interests (POIs) are displayed by black discs. **B** The displacement of the tagged dipole. x and y do overlap at 0 nm. **C** The signal measured at the different POI-locations as indicated in **A** for two periods of the ultrasonic wave. POI_4_ has the highest amplitude. **D** The absolute difference between co-elevation of optimal POI location (*θ*_POI_, i.e., point where highest signal amplitude is measured) and *θ*_dp_. **E, F** and **G** The signal strength at the optimal POI for POIs at the cortex (*R*_brain_), the scalp (*R*_scalp_) and with extra air-layer (*R*_air_), respectively. The gray horizontal line indicates the radial location of the POI.

For a dipole at *r*_dp_ = 20 mm and *θ*_dp_ = tan^−1^(1*/*2), the signal strength measurable at the scalp is shown in figure 2 (A). Here signal strength denotes the magnitude of the measured signal’s frequency component at *f*_us_ (i.e., |Ψ(*f* = 1 MHz) |). A hotspot between POI_2_ and POI_4_ can be observed. The signal strength does not monotonically decrease away from the hotspot. This is highlighted in figure 2 (C), where the signal at POI_5_, although closer to the hotspot, is lower than that at POIs 1 and 3. Furthermore, a clear oscillation of the signal can be seen at each POI. The oscillation is either in phase or antiphase with the ultrasonic displacement and has a period equal to 1/*f*_us_. For this setup, the maximal signal amplitude is only 0.1 pV.

For aforementioned dipole positions, the center of the hotspot is located in the yz-plane. The optimal polar angle (*θ*_POI_) depends both on *r*_dp_ and *θ*_dp_, as can be seen in figure 2 (D). When the dipole is close to the center, the optimal angle is between the dipole’s orientation and the displacement direction. At the center itself, this is the mean vector of both directions. On the other hand for a dipole closer to the outer surface, the optimal *θ*_POI_ is closer or equal to *θ*_dp_.

The maximal signal strengths measurable at the cortex, scalp or with an extra 5 mm thick air layer are shown in figures 2 (E), (F) and (G), respectively. As expected, the highest signals are measured with POIs at the cortex level (*R*_cortex_), while the lowest are measured with POIs in the air layer (*R*_air_, the dry electrode). The absolute maximum is 340.604 pV while the minimum is only 0.047 pV. For a cortical electrode, there is a three orders of magnitude difference between the lowest and highest measurable signal strengths. At the scalp (*R*_scalp_), this reduces to a difference of a factor 40. The lowest measured signal is half the one measured at the cortex. This while the maximum is more than a 100 times lower. The extra air layer reduces the maximum further with a factor 2, while the minimum is only reduced with 16%. For *θ*_dp_ = 0, introduction of the skull and scalp (and air) layer results in a signal reduction of 68% and 30% (11% and 7%) for a fixed distance of 22 mm and 70 mm with respect to the measuring electrodes, respectively. This slightly increases to 70% and 36% (11% and 9%) when the dipole orientation is perpendicular to the vibration direction. For all electrode setups, the minimum is for a dipole at the center with an orientation perpendicular to the displacement direction. The absolute maximum is found near the surface (*r*_dp_ = *R*_dp,max_) and for a *θ*_dp_ = 0. Consequently, maximal signal strength is obtained when the dipole orientation, displacement direction and POI position are perfectly aligned.

### B. Vibrational Interference

The applied ultrasonic waves propagate through the whole brain. A spatially distributed displacement field arises. In the focus, the displacement is maximal. The dipole located at this maximum is called the tagged region or dipole of interest (DOI). Although lower in amplitude, the dipoles outside the focus will vibrate as well. This will induce some interference on the signal of interest, as also these signals will be modulated onto *f*_us_. In this study, this type of interference is called vibrational interference. Moreover, the displacement field linearly scales with the transducer outputs. As such, only the displacement field profile itself is of importance for the vibrational interference generated by the surrounding dipoles. An synthetic displacement field is used, see eq. (12). Manipulation of the spatial constant *κ* allows for the investigation of the displacement field decay. The used PSD, eq. (7), for the surrounding dipoles consists of three cut-off frequencies with attenuation factors *α*_1,2_ = 2 and *α*_3_ = 10. This to remove static interference (see section III-C).

The results of the *κ* analysis are shown in figure 3. For a DOI at the center (see figure 3 (A)), reducing *κ* to 1 mm results in almost perfect reconstruction of the input signal (see figure 3 (B)). Also for *κ* = 5 mm, resemblance to the input signal is observed. In case of the cortex DOI (DOI at *R*_dp,max_, see figure 3 (D)), no and rather small resemblance between input signal and reconstructed is observed for *κ* = 5 and 1 mm, respectively. This is reflected in the RMSE as depicted in figures 3 (C) and (F). The RMSE is below 3% at almost all POIs for a deep DOI and *κ* = 1 mm. This increases to values between 20 and 40% for a *κ* = 5 mm. It increases above 60% for a cortex DOI (*κ* = 5 mm), with only values of 30% for *κ* = 1 mm. Also a difference in optimal POI position can be observed, for the deep DOI the optimal is POI_3_ and the worst is POI_5_. While the best POI for the cortex DOI is POI_2_ (equal to POI_4_). This being in agreement with the conclusion from section III-A. Inclusion of the artificial POIs, mPOI and mPSO, does not result in significant improvement. It is clear that even smaller *κ* is required for the signal of interest to be higher than the vibrational interference, in case of the cortex DOI.

**Fig. 3.**
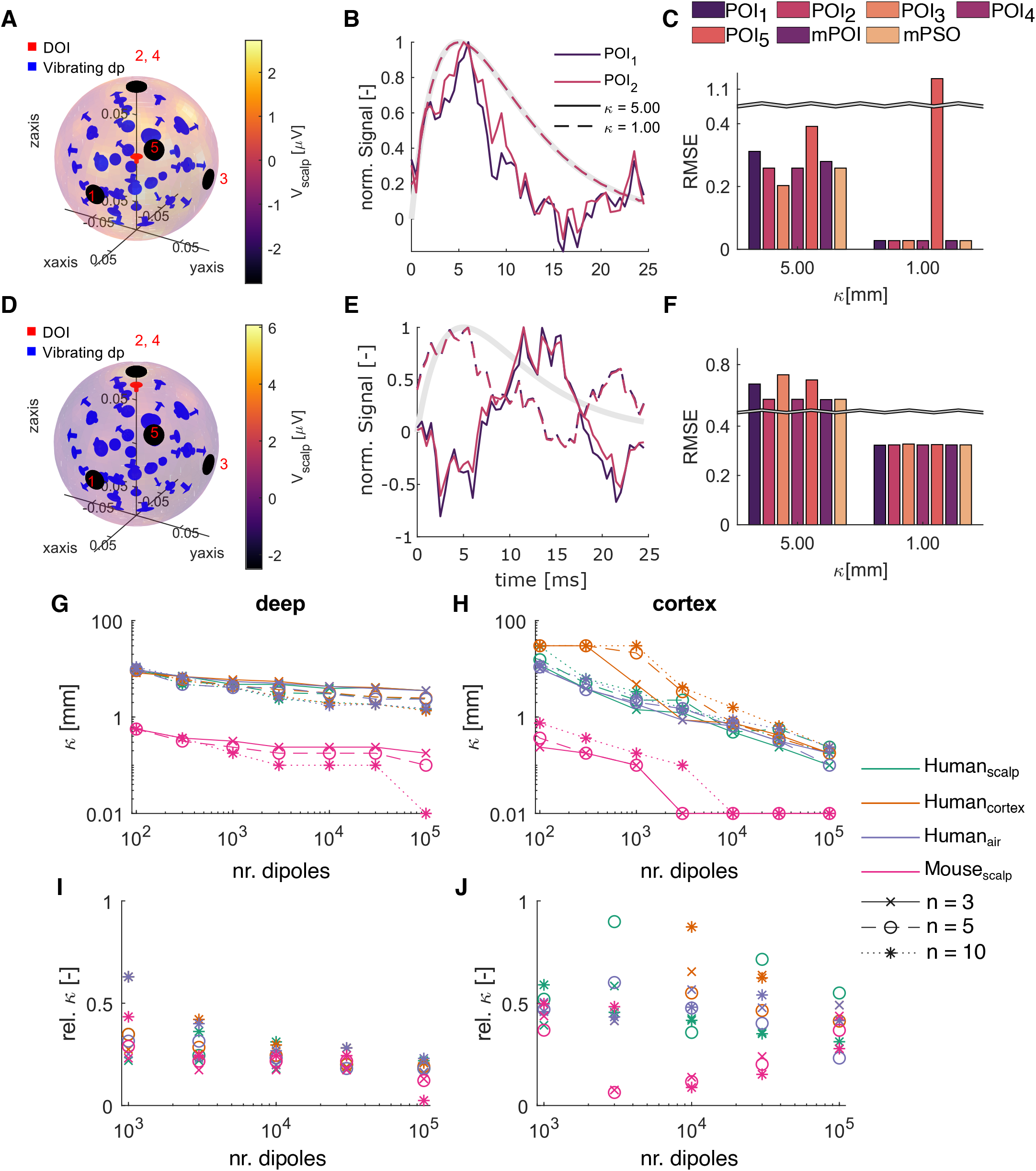
The effect of the displacement field’s spatial constant (*κ*) on vibrational interference and reconstructed signal quality. **A** Simulation setup for dipole of interest (DOI) at the center (deep) indicated by a red arrow. Blue arrows are the surrounding dipoles responsible for the vibrational interference. The POIs are indicated by black discs. The color code indicates the actual measured potential at a single time point. **B** The demodulated signal measured at indicated POIs for different spatial constants. The applied current intensity to the DOI is shown in gray. **C** The root mean square error (RMSE) of the demodulated signals measured at indicated POIs. mPOI, demodulation after the mean of the signals at the five POIs is taken first. mPSO, the mean of POIs with same order of magnitude before demodulation. **D** Simulation setup for DOI at *r*_dp_ = 65 mm (cortex). **E** and **F** Similar to **B** and **C**, respectively, but for DOI at the cortex. **G** and **H** The required spatial constant to get an RMSE of 5% on at least one POI for indicated models and number of dipole layers (*n*) with increasing number of surrounding dipoles, for a deep and cortex DOI, respectively. **I** and **J** The required relative spatial constant, i.e., *κ* divided by the distance between DOI and its nearest neighbor. In **B, C, E** and **F** The result shown is for 10^5^ dipoles and *n* = 3 layers, measured in the human scalp setup. To remove static interference, a PSD with three cut-off frequencies is used for the surrounding dipoles current intensity.

The required *κ* to get on at least one POI an RMSE ≤5% is shown in figures 3 (G) and (H), for a deep and cortex DOI, respectively. It can be noted that the measurement position, i.e. at the scalp, cortex or air, has little to no effect on the required *κ* for the deep DOI in the human head model. In case of the mouse model this is one order of magnitude lower. For a cortex DOI, in comparison with a deep DOI, a higher *κ* is acceptable for a relatively small number of dipoles. However, *κ* drops more quickly with increasing number of dipoles. The number of layers (*n*) has a clear influence in case of the deep DOI, as well. With a higher number of layers, a lower *κ* is required for all model setups. For the cortex dipole, this is overall opposite with a lower required *κ* for the lowest amount of layers.

A slightly decreasing and no clear trend can be observed for the relative *κ* shown in figure 3 (I) and (J), respectively. Here, relative *κ* (rel. *κ*) equals to *κ* divided by the distance between DOI and its nearest neighbor. For the deep dipole, the layer, and human versus mouse models differentiation disappears. From these relative *κ* plots, it is clear that the most important factor defining the required spatial constant is the distance between the DOI and its nearest neighbor. The mouse model is smaller. Therefore, for a same number of dipoles, it is more densely packed than the human models. The difference, for a changing number of dipole layers, between the deep and cortex DOI plots can be explained by this, as well. The *n* layers are distributed equidistant from 0 to *R*_dp,max_ (0 excluded). In case of the deep DOI and for a higher *n*, the closest neighbor is thus much closer than for a lower *n*. As a result, the required *κ* decreases with *n* for a deep dipole. In contrast, for a cortex dipole, the *κ*-threshold tends to increase with *n*, because the dipoles are uniformly distributed onto these layers. For a fixed number of dipoles, each layer will thus be occupied more in case of low layer count, reducing the distance between the DOI and the nearest neighbor on the same layer. The sudden drop between 10^3^ and 10^4^ dipoles indicates the shift from nearest neighbor on another layer to the same layer. The strong decreasing trend for a cortex dipole is thus explained by the decreasing distance to the nearest neighbor at the same layer. This while in case of the deep DOI (figure 3 (G)), the slight decrease is due to combined contribution of all vibrating dipoles because the distance to the nearest neighbor, located at the next layer, is fixed.

### C. Static Interference

A second type of interference originates from the electrical activity in the brain itself. In this study this is called the static interference. Because the EEG energy at MHz-range is not known, the applied current intensities to the dipoles are based on a power-law power spectral density, see eq. (7). Unlike in the previous section, the PSD only consist of two cut-off frequencies. For *α*_2_ = 2, this results in a drop of 110 dB at *f* = 1 MHz, with respect to DC (f = 0 Hz). To minimize the vibrational interference, half of the required *κ* to achieve an RMSE of 5% with 10^5^ dipoles is used. The values are summarized in table II. Increasing the displacement amplitude (*A*_max_) increases the signal-to-noise ratio with respect to the static interference. The results of the *A*_max_ analysis are shown in figure 4.

**Table II.**
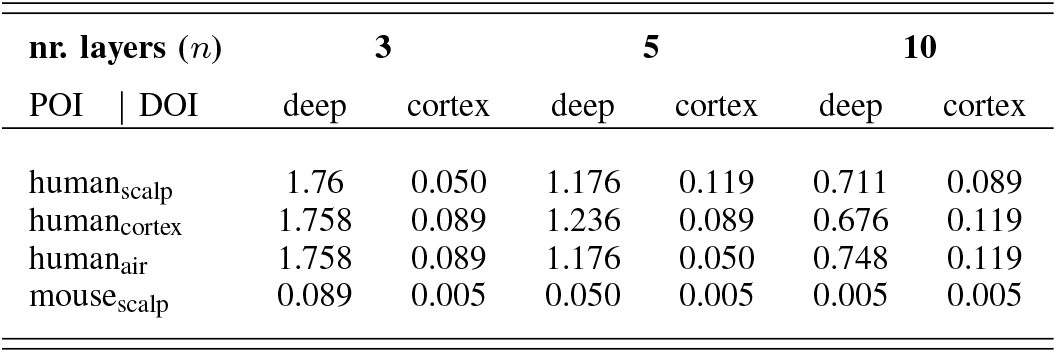
Used spatial constants (*κ*) in displacement amplitude simulations. Half of *κ* to get rmse of 5% for 10^5^ dipoles.

**Fig. 4.**
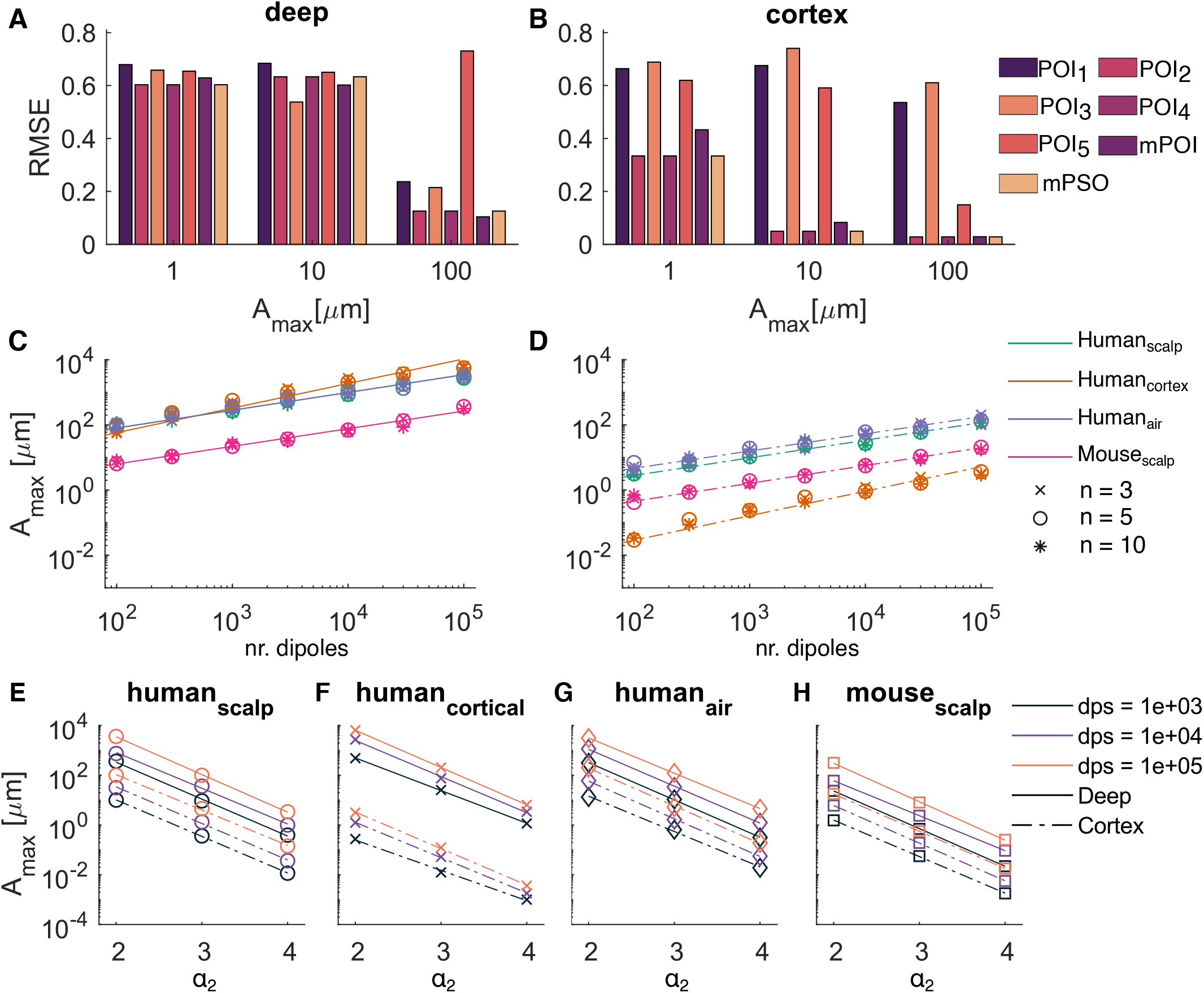
Analysis of the necessary displacement amplitude (A_max_) to overcome the static interference. **A** and **B** The root mean square error (RMSE) between target and demodulated signal measured at indicated POIs with different A_max_, for deep and cortex DOI, respectively (*n* = 3 layers and nr. of dipoles dps = 10^5^, measured in human scalp setup). mPOI, demodulation after the mean of the signals at the five POIs is taken first. mPSO, the mean of POIs with same order of magnitude before demodulation. **C** The required A_max_ to obtain an RMSE of 5% on at least one POI for indicated models and number of dipole layers (*n*) with increasing number of surrounding dipoles. Results are for a deep DOI. **D** DOI at the cortex. The plotted lines indicate the linear regression for each model and POI position combination. **E, F, G** and **H** Effect of the second attenuation coefficient (*α*_2_) on the required A_max_ (*n* = 3 dipole layers). The used model and electrode position is shown in the title.

The RMSE of the reconstructed signals obtained at the different POIs for the human scalp setup, are shown in figures 4 (A) and (B). As expected, with increasing *A*_max_, the RMSE drops at almost all POI positions. POI_2_ is the best location, both for deep and cortex DOI. The necessary *A*_max_ to get an RMSE *<* 10% is 1 to 2 orders of magnitude lower for the cortex than the deep DOI.

The *A*_max_ to obtain an RMSE = 5%, for a deep and cortex DOI, are shown in figure 4 (C) and (D), respectively. For each model setup, the required *A*_max_ increases linearly on the log-log scale, with increasing number of dipoles, i.e., log^10^ *A*_max_ = *a* log^10^(nr. dipoles) + *b*. The different model setups have similar slopes, around 0.54. Except the cortex measurements, these have a slope around 0.75. Interesting is the different order of model setups between the deep and cortex DOI. In case of the former, the highest displacement amplitude is required for a human cortex setup, while this requires the lowest amplitude when the DOI is at the cortex. The extra air layer has negligible effect for the deep DOI with an intercept difference of only 0.0021. On the other hand, there is a significant effect when the DOI is at *r*_dp_ = *R*_dp,max_, leading to an intercept difference of 0.2439. Moreover, there is a 1 (2) order(s) of magnitude difference between the mouse and human model in case of the deep (cortex) DOI. Finally, It can be seen that the number of layers (*n*) has no effect on *A*^max^.

The static interference is strongly governed by *α*_2_. As the real value is unknown, the effect of this parameter was further investigated. As shown in figures 4 (E-H), the required *A*_max_, to have an RMSE = 5%, decreases with increasing *α*_2_ for each model setup and fixed number of dipoles. Moreover, a similar slope around -1.5 is observed in case of all model setups and DOI locations. From eq. (7), the relationship between *α*_2_ and the level of dipole moment power *L*_P_ at *f*_us_ compared to DC is roughly: *L*_P_ [dB] = 10 log^10^ (PSD(*f* = 1 MHz)*/*PSD(*f* = 0 MHz)) ≈ −30*α*_2_ −50. Consequently, 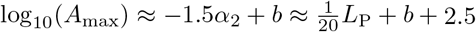, where *b* depends on nr. dipoles, DOI position and model setup. The signal strength increases linearly with *A*_max_ (see eq. (3)) Therefore, the power increases with 20 log^10^ *A*_max_, explaining the obtained relationship.

For the technique to be safe, the FDA limits (see section II-F) may not be exceeded. The minimal *α*_2_ to have an RMSE *<* 5% for a displacement amplitude of 50 nm is determined via interpolation and summarized in table III. For this *A*_max_, the corresponding MI and *I*_pa_ are 0.45 and 7.4 W/cm^2^, respectively. Consequently, a maximal dc of 9.74% is allowed in order to meet the *I*_*ta*_ limit for our pulse active time of 25 ms. The corresponding power *L*_P_ (in dB) is shown as well. It is clear that for a deeper DOI the static interference should be much lower.

**Table III.** Summary of the second attenuation coefficient needed to have an Rmse *<* 5% with a displacement amplitude of 50 nm. *l*_p_ denotes the level of dipole moment power *l*_p_[*db*] = 10 log_10_ (psd(*f* = 1 mhz)*/*psd(*f* = 0 mhz)).

| nr. dipoles |  | 1000 |  | 10000 |  | 100000 |  |
| --- | --- | --- | --- | --- | --- | --- | --- |
| DOI | POI | $\alpha_2$ | $-L_P$ | $\alpha_2$ | $-L_P$ | $\alpha_2$ | $-L_P$ |
| deep | human<sub>scalp</sub> | 4.58 | 184 | 4.93 | 194 | 5.20 | 202 |
|  | human<sub>cortex</sub> | 5.05 | 198 | 5.24 | 203 | 5.41 | 209 |
|  | human<sub>air</sub> | 4.54 | 183 | 4.93 | 194 | 5.38 | 208 |
|  | mouse<sub>scalp</sub> | 3.76 | 159 | 4.19 | 172 | 4.45 | 180 |
| cortex | human<sub>scalp</sub> | 3.57 | 154 | 3.93 | 164 | 4.34 | 177 |
|  | human<sub>cortex</sub> | 2.57 | 123 | 2.99 | 136 | 3.24 | 144 |
|  | human<sub>air</sub> | 3.73 | 158 | 4.02 | 167 | 4.41 | 179 |
|  | mouse<sub>scalp</sub> | 3.02 | 137 | 3.38 | 148 | 3.70 | 157 |

### D. Reconstruction accuracy and dipole current waveform

Finally, the effect of the chosen target parameters is addressed. The results above were for a target RMSE of 5%. Ideally, a perfect reconstruction of the target’s current intensity profile could be obtained. Due to the aforementioned interferences, this can only be achieved with an infinitesimal small *κ* and infinitely high *A*_max_. Capturing the general trend on the other hand can be informative as well. To investigate the effect of the imposed reconstruction accuracy, the required *κ* and *A*_max_ for 10^5^ dipoles are determined for an RMSE of 10%. Also, the current intensity profile of the DOI is switched from an alpha-function to an alpha-train (see eq (11) and figure 1 (D)). The relative change in parameters are shown in figure 5 (i.e., the required *κ* (*A*_max_) for the new target settings, either altered input current intensity (alpha-train) or altered RMSE target value (10%), divided by the required parameter value for the alpha-function intensity profile and RMSE = 5%). The change of target function is expected to give values of 1 for both 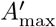 and *κ*^*′*^. In case of some simulation setups, there is a large deviation from this value. From figure 3 (B), it was already concluded that the RMSE is a very strict metric for the alpha-function input. Moreover, due to the interferences, the reconstructed signal will be noisy. This can potentially favor highly time-varying signals such as the alpha-train.

**Fig. 5.**
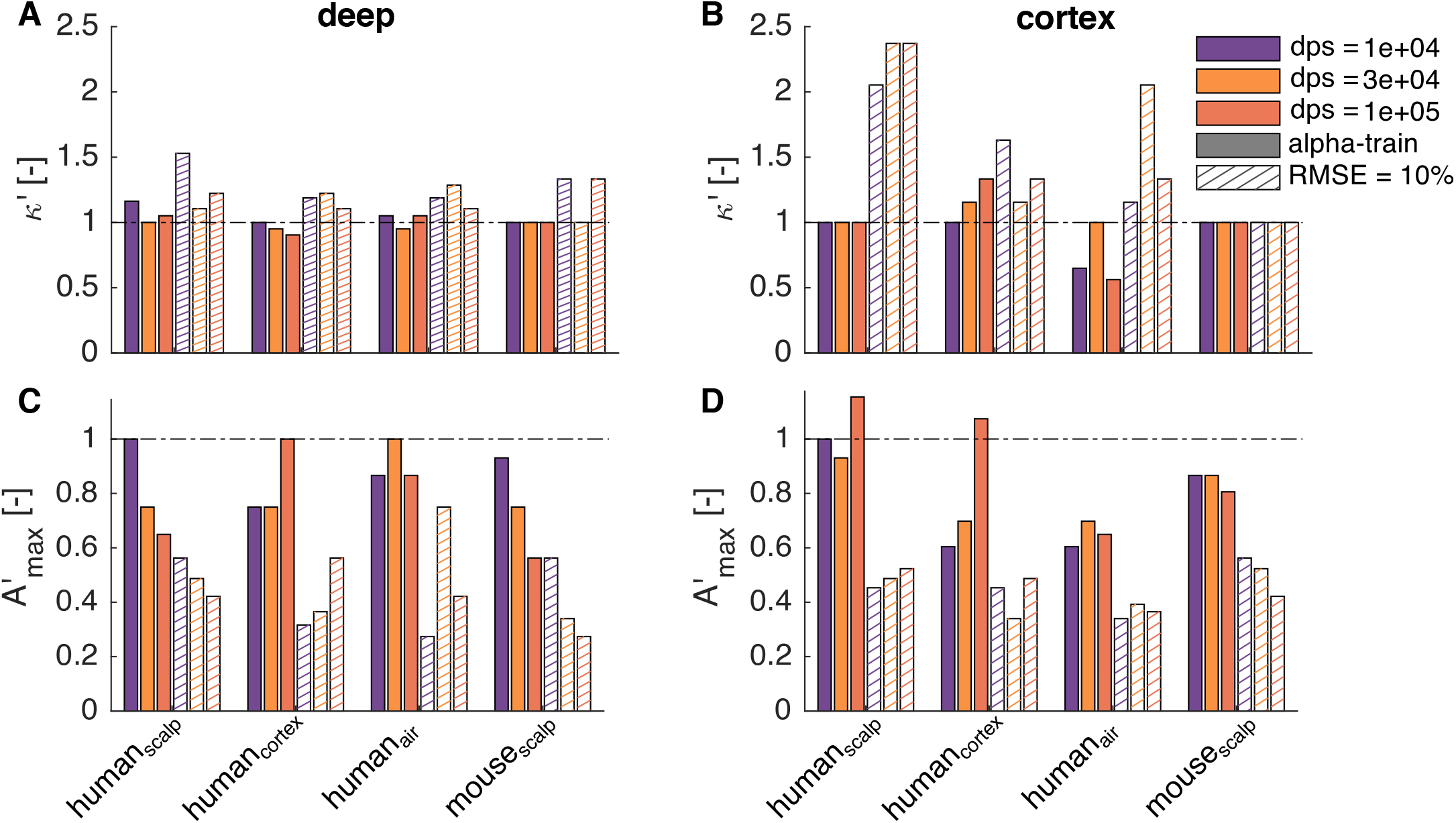
The effect of an altered intensity current (alpha-train) and changed cut-off RMSE level (10%). **A** The relative change in spatial constant *κ* with respect to simulation with original target settings, i.e., alpha-fun and RMSE of 5%, for the different models. The dipole of interest (DOI) is at *r*_dp_ = 0 mm. The results shown are for different number of surrounding dipoles. **B** a DOI at *r*_dp_ = 65mm (cortex). **C** and **D** The relative change in required displacement amplitude (A_max_) with respect to simulation with original target settings, i.e., alpha-fun and RMSE of 5%.

For a target RMSE of 10%, an increase in *κ* and a decrease in *A*_max_ is expected. Overall, this is observed although for some setups less pronounced. In case of the deep DOI, only small changes (*<* 1.5) in *κ* are observed. These go up to 2.5 for the cortex dipole. For almost all setups and DOI locations, *A*_max_ is halved or lower. It is clear that the opposed restrictions can have significant effect but do not result in differences in several orders of magnitude, which is more of interest in this simulation study.

## IV. Discussion

The results in this study showed that it is possible to reconstruct the input intensity profile current of a mechanically vibrating dipole from the frequency content near the ultrasonic frequency. In the used model, the head is represented as a set of concentric spheres and the signal generators as dipoles. An optimal signal strength is obtained when dipole orientation, vibration direction and POI are perfectly aligned. When perpendicular, this is minimal. Inclusion of a skull and scalp layer causes a reduction in maximal signal strength of two orders of magnitude. An extra air layer causes an extra reduction of a factor 2. The smallest signals only decrease by a factor two and 16% by including a skull-scalp and air layer, respectively. It is clear that that there is a large signal reduction due to the extra distance between source and receiver. The introduction of extra layers account for large additional losses (both propagation and conductive). Conductive loss is more prominent with the introduction of the skull and scalp tissues. The signal strengths are, however, small. For a fixed dipole moment of 5 nAm and a vibration amplitude within FDA limits, the strength is in the order of pV.

Two interference types are identified and investigated. First, the vibrational interference which originates from a vibrating region excluding the dipole of interest. Second, the static interference that comes from the electrical activity of the regions itself. Concerning the vibrational interference, it was found that lower spatial constants are needed to achieve accurate reconstructed signals (i.e., RMSE*<*5%) with increasing number of dipoles. The *κ* in the mouse model is on average an order of magnitude lower than that in the human head model setups. Although opposite for the deep and cortex DOI, there is also a clear effect of the number of layers (*n*) modeled. As aforementioned (section III-B), these three aspects can be explained by taking into account the distance between DOI and nearest neighbor, as also indicated by rel. *κ*. It should be noted that all dipoles vibrate in phase. Consequently, the current vibrational interference reflects the upper limit. For the cortex DOI, also an effect of the electrode position is observed. This is due to the strong non-linear dependence between dipole vibration direction, orientation and POI position as is shown in figure 2. This is more pronounced if the dipole is closer to the POI than further away. Meaning that the contributions of neighboring dipoles are relatively lower for the cortex setup than for the scalp and air setups (see cortex versus scalp or air POI in figure 3 (G) and (H)).

The difference in *A*_max_ between the deep and cortex DOI is completely explained by the signal drop due to increasing distance between source and observer (see figure 2). This holds also for the deep mouse scalp setup. The reason for the mouse scalp setup to require a larger *A*_max_ than human cortex is attributable to the relative distance difference between the DOI and the surrounding dipoles to the POI. Finally, the increased slope of the required *A*_max_ with increasing number of dipoles for human cortex measurement can be explained by the increasing density of dipoles near the POI. This due to the 1*/r*^*i*^ dependence as seen eq. (S.3). When the distance is small (cortical electrode), a small change has relatively a higher impact than when initial distance is larger (scalp electrode). Consequently, a steeper slope is expected for the scalp versus air setup but this is below the simulation setup’s accuracy limits.

Focusing an ultrasonic beam is complicated by the distortion imposed by crossing the skull [38]. Although, adaptive focusing techniques exist, determining the field for each setup is too tedious and therefore out of the scope of this study. Moreover, a field induced by a transducer array is not monotonically decreasing but contains sidelobes. The sidelobes’ characteristics strongly depend on the transducer setup [39]. To avoid dipoles to be located at the trough of a sidelobe, a monotonically exponential decaying field was selected (see eq. (12)). With its spatial constant, the selected field distribution allows for systematic investigation, capturing the essential spatial decay from the ultrasonic focus, which could be used as guideline for transducer development in the future. Moreover, the spatial constant is strongly correlated to the spatial resolution that be obtained by acousto-electrophysiological neuroimaging, however, limited by the ultrasonic field. As discussed above, the closer the nearest neighbor, the lower the required *κ*. Or in other words, for a fixed *κ* the distance to the nearest neighbor is defined, this being the spatial resolution. For the cortex DOI in the human (mouse) model, this is 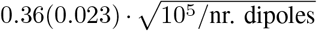 mm. In case of the deep DOI, this is 6.5 mm (0.41 mm).

The dipole is an anatomically constraint representation of current generators, a model that is typically used in ESI. Murakami 2015 showed that, in the brain, the dipole density has a maximum value between 1-2 nAm/mm^2^ across different species. The dipole moment itself can vary over 1-3 orders of magnitude depending on the volume of active tissue. A cortical column comprises around 10^5^ cells/mm^2^. Consequently, with a single neuron’s dipole moment between 0.1-1 pAm [40] this results in a synchronously active fraction of 1-20% [34].

For the investigated method to be safe, the FDA limits cannot be exceeded. The important parameter here is *A*_max_. The required *A*_max_ is determined by the static interference and thermal or instrumental noise. The latter is not included in this study as it is defined by the recording device (EEG-instrumentation). As the frequency content of the brain at US-frequency range is not known, the input current intensities were drawn from a power-law power spectral density profile. This has been studied in literature as this 1*/f*^*α*^-like power spectrum is observed at many spatio-temporal scales. Power-laws with *α* between 0-4 have been reported depending on the measured scale for frequencies below 1 kHz [41]–[43]. Dipole moment calculation were performed on morphologically accurate neuron models [44], [45] to gain more insight in the energy content of realistic current dipoles for f *>* 1 kHz [46]. Our calculations showed fitted power law coefficients of 4 to 5 for f *ϵ* [1, 25] kHz (see supplementary section IV). This corresponds to an *α*_2_-value of 3 to 4. Although not conclusive, it justifies the tested PSD profiles but further investigation is necessary. The results shown in table III showed that if the level of dipole moment power (*L*_P_) is -210 dB at 1 MHz, good signal reconstruction (RMSE = 5%) can be obtained for all tested model setups with only a displacement amplitude of 50 nm. This amplitude is well within FDA limits, if a dc of 9.74% is used with our pulse active time of 25 ms. As expected, the *L*_P_ correlates with the number of dipoles. These dipoles represent a small volume of neurons. Therefore, the number of dipoles and their strength *I*_max_ is limited, depending on the represented volume. Although the static interference level depends on the number of dipoles in this simulation setup, the static interference will be fixed in the brain. On the other hand, the strength of the tagged region decreases with increasing spatial resolution as the represented volume decreases. Consequently, the signal-to-noise ratio still drops with increasing nr. dipoles and the same effect on the required *A*_max_ is thus expected. Therefore, because the number of dipoles dictates the spatial resolution, retrospectively, if the static interference is known, a decision can be made whether or not and at which resolution the technique will be safe, within FDA limits.

### A. The Acousto-Electric Effect

Witte et al. experimentally validated that indeed the electrophysiological activity gets modulated on the harmonic frequency when selectively probed with ultrasound. This was shown in numerous experiments, e.g., in a rabbit heart [28], [47], lobster nerve [25], bath with 0.9% NaCl solution [24], [27] and a human head phantom [29], [30]. Broadband-EEG like signals can be retrieved and current sources localized with high spatial accuracy. The latter is defined by the US focal zone, being only a couple of millimeters, not only for superficial but also for deep regions [30]. By scanning with the ultrasonic beam, a 4D image can be made with unprecedented spatiotemporal resolution. Results of this group and others are summarized in Zhang et al. (2022) [26]. The postulated underlying mechanism, the acousto-electric effect, i.e., a pressure induced change in resistivity, differs from the one hypothesized here. The mathematical formulation of modulation due to the AE is as follows:

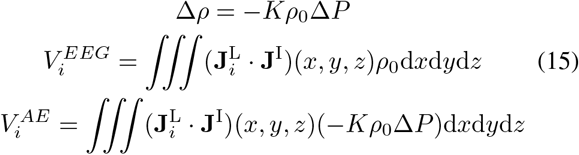

with *ρ*_0_ the direct current resistivity, *K* the acousto-electric interaction constant, Δ*P* the ultrasound pressure field, and 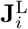 and **J**^I^ the lead field of lead i and current source density field, respectively. For the derivation we refer to Olafsson et al. (2008) [24]. In the literature on the acousto-electric effect, the potential term 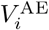 is caused by ultrasound-induced oscillations of the electrical resistivity Δ*ρ*, while the lead field 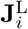 and the current source **J**^I^ are implicitly assumed to be undisturbed by the pressure field. However, we argue that the AE-induced resistivity oscillations will cause direct changes to the current density fields (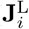 and **J**^I^). Conversely, in this study, modulation of the DOI current on the ultrasonic sine occurs due to the relative motion of the dipole of interest. Consequently, both the acousto-electric effect and the vibration of dipoles (or equivalently, the oscillation of current sources **J**^I^ and lead fields 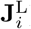) can be interpreted as complementary tentative underlying mechanisms of acousto-electrophysiological neuroimaging. Furthermore, the results of this study clearly show that the latter mechanism (vibrating dipoles) also suffices to modulate the endogenous signal onto the ultrasonic frequency.

Wang et al. (2011) [27] reported a 27 nV/mA peak signal strength with 500 kPa pressure, and 5 mm distance between current source and sink, measured at 5 mm from current source. This equals to 10.8 pV*/*(nAm *·* MPa). Performing an equivalent simulation, i.e., dipole at [0, 0, 0], orientation [0, 1, 0], vibration direction [0, 0, 1], *d* = 5 mm and electrode at [0, 7.50, 0] mm, results in 6.1 10^−6^ pV*/*(nAm *·* MPa). However, our results showed a strong nonlinear dependence on the alignment between the dipole, displacement direction and electrode position. Subsequently, a 1% misalignment (electrode at [0, 7.50, 0.075] mm) gives 1.8 pV*/*(nAm *·* MPa). It should be noted, that the dipole vibrates as a whole. Moreover, *d* only affects the dipole moment as no distinction between current source and sink is made (see eq. (S.3)). Taken this into account, still, a comparable strength is expected for a vibrating current dipole. A maximum of 91.0 pV*/*(nAm *·* MPa) is obtained with an electrode at the optimal POI location [0, 5.3, 5.3] mm.

### B. Limitations and Future Work

As already mentioned in section III-D, the used RMSE metric is prone to randomness in the results. It is very strict in the sense that only low values are returned when almost perfect profile match is obtained. As visible in figure 3 (B), already a clear peak is observed for *κ* = 5 mm, while only a minimum RMSE of 20% was calculated. Consequently, more favorable spatial constants or displacement amplitudes than the ones determined for RMSE = 5% could already give feasible reconstructions, for instance when scanning for hyperactive regions, like in epilepsy. Here, the dipole moments will be 1-2 orders of magnitude higher as well, increasing the signal strength with the same amount [34]. On the other hand, as depicted in figure 5, sometimes no clear trends can be observed. This is due to the randomness introduced in the simulations (e.g., the current intensity profile and strength, and dipole positions). Simulations could be repeated for different random seeds. However, it was deemed not feasible due to the need for large computational resources and the expected change only being in same order of magnitude.

There was opted not to use frequency dependent electrical properties for the different tissues. This is because the frequency dependence for the sub-MHz range is not clearly established [48]. Moreover, the small changes do not outweigh the big increase in computation time. A single test was performed with the frequency-dependent values obtain from [33], which resulted in a 20% increase of the signal strength received from a single dipole. Also, the electrical properties were set to be homogeneous, confined to each sphere, and isotropic. However, experiments have shown those to be inhomogeneous and anisotropic [9]. This will clearly impact the signal strength. However, the effect was deemed to be inferior to the spherical approximation with respect to the correct morphological shape. Moreover, the in-vivo values are still under debate [21]. While this impacts ESI greatly, with possible localization errors up to several centimeters, here this will only impact the signal strength without affecting the achievable resolution.

In this study the AE-effect is not taken into account (i.e., the conductivity is assumed to be non-oscillating). In future research, the acousto-electric effect could be included. Moreover, further research is necessary to determine if one of the proposed underlying mechanisms of AENI (vibrating current sources and the AE) is dominant in the brain for given conditions (e.g., waveform, transducer placement, dipole location). The brain regions are modeled as rigid dipoles. As such, the current source and sink vibrate in phase. The contribution of stretching and rotation could be further investigated. Also, the dipole assumption itself compared to discrete monopoles or a multipole expansion could be of interest in future work. The ultrasonic frequency was kept constant throughout the whole study. Lowering the frequency will both lower the MI and the *I*_pa_ for a fixed displacement amplitude, but this will increase the static noise.

In future work, the added value of modulated focused ultrasound [49] can be investigated with the goal of reducing the MI. This would be beneficial, if the acoustic particle velocity amplitude is proportional to the beat frequency *f*_b_ (*v*_us_ ≈ 2*πAf*_b_) due to the low-pass filter property of the constitutive equation of the viscoelastic brain. In this case, the mechanical index is 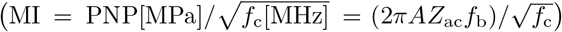. In other words, a small ratio of beat to carrier frequency 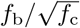, results in a smaller mechanical index. Finally, a limit on possible neuromodulation should be considered, as well. Indeed, the goal of AENI is to record endogenous activity by ultrasonically vibrating a region of interest, but ultrasound has been demonstrated to modulate brain activity in animals and humans [50], [51]. In future work, computational modeling of ultrasonic neuromodulation can be used to determine the region of the parameter space in which AENI is feasible without significant modulation of the endogenous activity [52]–[54]. Although a single 25 ms ultrasonic pulse was adopted in this study, signal reconstruction is still possible with pulsed ultrasonic fields with a signal period of 25 ms. In the current literature, relatively large duty cycles are adopted in order to achieve neuromodulation [55]. Moreover, an exponential increase of neuromodulation thresholds is observed with lowering the duty cycle [56], while only a linear relationship is expected concerning the signal-to-noise ratio with respect to the static interference.

With pulsed fUS, a rise in recorded activity at the pulse repetition frequency (prf) was also observed in [57]. It is expected that the measured signal at the prf is either the signal induced by neuromodulation (as stated by the authors of [57]) [56], [58] or by hearing confounds [59], [60]. It is hypothesized in [57] that also endogenous activity of the targeted area could be modulated on the recorded signal at the pulse repetition frequency. However, based on the mechanism described above, the relative signal strength at *f* = prf with respect to the strength at *f*_us_ should be sinc((prf − *f*_us_)*/*(prf*/*dc)). Using the values reported in Darvas et al. (2016) [57] this results in ≈ 5 *·* 10^−5^, implying that demodulation of endogenous activity at the repetition frequency is unlikely, at least for the vibrating dipole mechanism of AENI.

Feasibility of acousto-electrophysiological neuroimaging will depend on the currently unknown biophysiological activity at ultrasonic frequencies. Unlike with high-density EEG, accurate reconstruction of a tagged region’s signal is possible, with just one electrode (and one reference). Moreover, no MRI or complex ESI algorithm is necessary to solve the ill-posed inverse problem. On the other hand 4D imaging with UCSDI currently takes up several hours in experimental setups. This could probably be dramatically improved with modern scanners. Additionally, B-mode ultrasound and normal EEG could be coregistered with AENI [27]. Finally, spatially encoding the brain with different ultrasonic frequencies could be an interesting path to investigate in the future.

## V. Conclusion

This feasibility study showed that mechanical vibration, introduced by an ultrasonic field, modulates the endogenous signals onto the ultrasonic frequency. In this spherical representation of the head where the active brain is discretized into a set of dipoles, the signal strength strongly depends on the alignment between dipole moment, displacement direction and point of measurement. Inclusion of extra layers results in a signal reduction, attributable to the extra distance between source and receiver and conductive losses. For a displacement amplitude of 10 nm and a dipole moment of 5 nAm the signal strengths are low, with a maximal signal strength of 341.60, 2.36 and 1.24 pV that can be measured with a cortical, wet and dry scalp electrode, respectively. For a dipole at the center, the strengths are below 0.16 pV. Accurate reconstruction of a tagged region’s activity can be obtained if two interference sources are overcome. The vibrational interference originates from other vibrating regions. This can be decreased by decreasing the spatial constant of the ultrasonic field. It was shown that the dominant factor is the distance to the nearest neighboring dipole. Therefore, the spatial resolution is strongly correlated to this spatial constant. The static interference comes from the endogenous activity of non-vibrating brain regions self at the ultrasonic frequency. The signal-to-noise ratio increases with increasing displacement amplitude. This is, however, limited by the mechanical index and average pulse intensity limits set by the FDA. Log-log relationships are observed between the required displacement amplitude and the power of the static interference at *f*_us_, and the number of dipoles. For a deep region of interest, dry and wet electrodes deliver similar and best results. An accurate signal reconstruction (RMSE *<* 5%) can be obtained if the level of dipole moment power ≈ − 154 − 10 log_10_(nr.dipoles) dB. This under safe conditions with only 50 nm displacement amplitude. For the cortex region, cortical electrodes give the best result, with a required level of dipole moment of ≈ − 94 − 10 log_10_(nr.dipoles) dB. With the mouse model, lower vibration amplitudes are required for detection, but a spatial constant in the order of 10 *μ*m is required. Depending on the spatial constant, resolutions up to millimeter could theoretically be achieved in humans but will completely depend on the ultrasonic field.

## Acknowledgment

R. Schoeters is a PhD Fellow of the FWO-V (FR) (Research Foundation Flanders, Belgium). T. Tarnaud is a postdoctoral fellow of the FWO-V.

We would like to thank Xavier Rottenberg and Peter Peumans of IMEC Belgium, for their insights, shared knowledge and interesting discussions.

This work was carried out using the Supercomputer Infrastructure (STEVIN) at Ghent University, funded by Ghent University, the Flemish Supercomputer Center (VSC), the Hercules Foundation and the Flemish Government department EWI.

## Supplementary Info

### S.1. The LiÉnard-Wiechert Fields

The Liénard-Wiechert field, which is the time varying electromagnetic field for a point charge (q) at position (**r**_**s**_) in arbitrary motion, is given by:

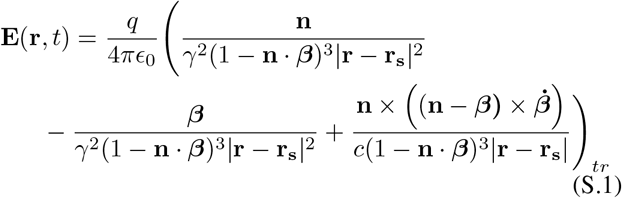

with *ϵ*_0_ the vacuum permittivity and *c* the speed of light. **n**(*t*) = (**r r**_**s**_(*t*))*/* |**r** − **r**_**s**_(*t*)|, ***β***(*t*) = **v**(*t*)*/c*, where **v**(*t*) is the source’s velocity. 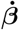 is the charge’s acceleration with respect to *c* (i.e., **a**(*t*)*/c*, with **a**(*t*) = d**v**(*t*)*/*d*t*), *γ*(*t*) is the Lorentz factor 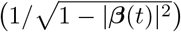 and *t*_*r*_ = *t* − |**r** − **r**_**s**_| */c* is the time retardation.

It can be divided into five components:

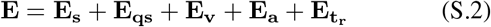

where

1. The static field: **E**_**s**_ **= E** when **v** = **0** and **a** = **0**
2. The quasi-static field: **E**_**qs**_ = **E** − **E**_**s**_ when *v << c* and 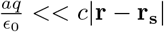
3. Relativistic effects due to velocity (Doppler): **E**_**v**_ = **E** − **E**_**s**_ − **E**_**qs**_ when 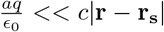
4. Relativistic effects due to acceleration (Electromagnetic Radiation): **E**_**a**_ = **E** − **E**_**s**_ − **E**_**qs**_ when *v << c*
5. Relativistic effects due to finite propagation speed: **E**_**tr**_ = **E** − **E**_**s**_ − **E**_**qs**_ − **E**_**v**_ − **E**_**a**_ when *t* ≠ *t*_*r*_

For a oscillating movement in the x-direction (*A* sin(*ωt*)**e**_*x*_) and an observer on the y-axis or x-axis, the dominant term of th Taylor expansion for A→0 is given in table IV. The quasi static contribution dominates if 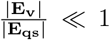 and 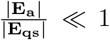.

A critical frequency can be found form 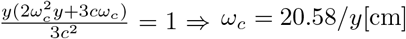 GHz. For an overestimation of y = 20 cm, *ω*_*c*_ = 1.029 GHz. Thus, for ultrasonic frequencies in the MHz range the quasi-static contributions dominate (*ω*_*us*_ ≪ *ω*_*c*_).

**Table IV.**
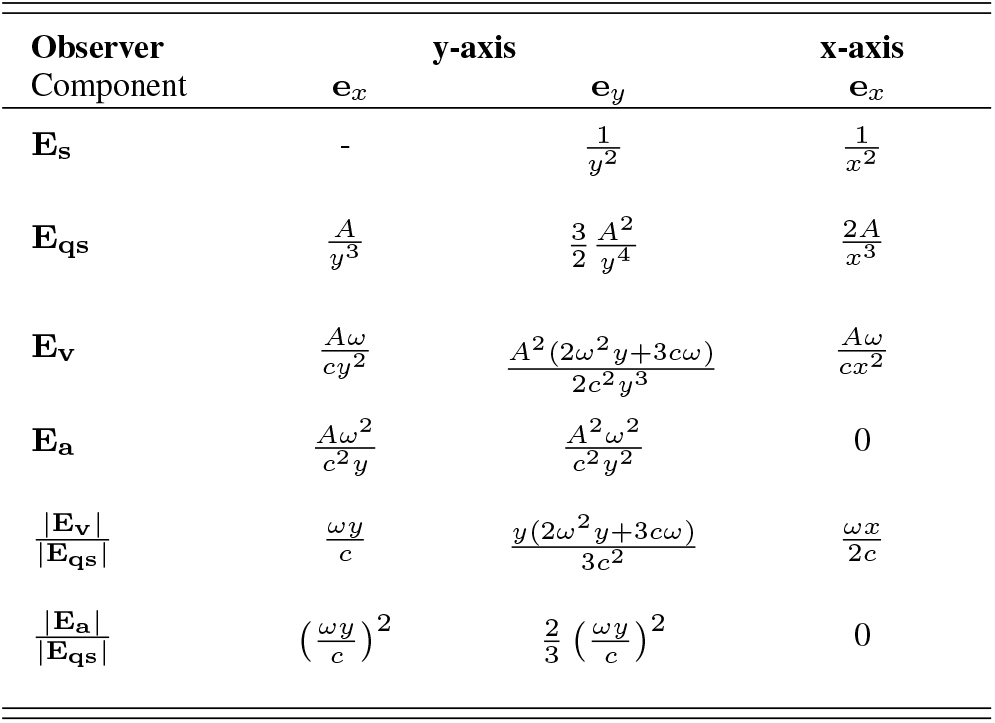
Dominant term of taylor expansion (a→0) of each component to the electromagnetic filed. Motion is in x-direction: *a* sin(*ωt*)**e**_*x*_. All values divided by *q/*4*πϵ*_0_

### S.2. SOLUTION TO GENERAL POISSON EQUATION SPHERICAL 4-LAYER HEAD MODEL

A semi-analytical solution can be obtained by solving the general Poisson equation in spherical coordinates. For a current dipole with moment *I*_max_*f* (*t*)**d**_dp_ on the z-axis (see manuscript figure 1 (C)), the induced electric potential *V*_*l*_ in volume *l* for an arbitrary point on the XZ-plane can be determined via:

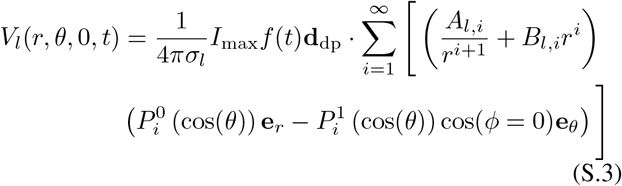

where *σ*_*l*_ is the conductivity of volume *l* (manuscript figure 1 (B)). *r, θ* and *ϕ* are spherical coordinates. The terms 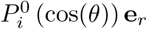 and 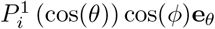 correspond to radial and tangential dipole, respectively, with 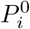 and 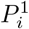 the associated Legendre polynomial of zeroth and first order [31], [32]. The Condon-Shortley phase is included in calculating the associated Legendre polynomials, explaining the minus sign between radial and tangential dipole terms (unlike in Arthur (1970) [31]). Values for *A*_*l,i*_ and *B*_*l,i*_ can be determined under following boundary conditions, i.e., the potential needs to be continues (1.) with reference at infinity (3.) and there is conservation of charge at the interface (2.):

1. *V*_*l*_ = *V*_*l*+1_
2. 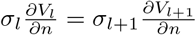
3. *V*_*l*_(*r*) → 0 for *r* → ∞

Due to 3., *B*_4_ has to be zero. *A*_1_ is defined by the unbound solution. This gives *A*_1_ = *b*^*i*−1^*p*, where *b* = |**r**_dp_| and *p* = |**d**|. Next, 1. provides a set of 3 equations:

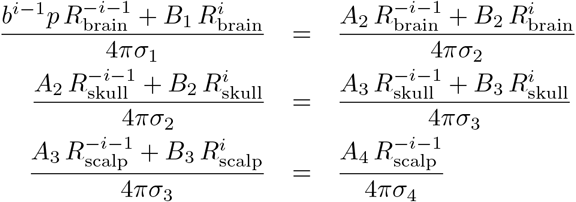

Finally, the last set of equations is given by 2.:

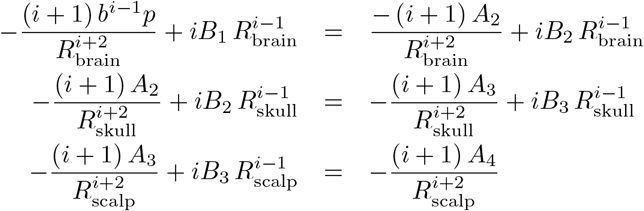

Consequently, the set of equations is defined and a solution can be found. The solution was obtained in Maple 2021.

### S.3. Time-Dependent Dipole Moment

In the model, dipoles are used to model the endogenous activity sources. Due to the interest in measured signal content at the ultrasonic frequency (*f*_us_), an appropriate dipole’s current intensity profile needs to be chosen to correctly capture the static interference (see manuscript section III-C). The simulations are solved with an sampling frequency *F*_*s*_ = 20 *f*_us_. To avoid distortion of the frequency content due to linear interpolation in the time domain, the time dependent signals are derived from a power spectral density profile (PSD). The formulation of the PSD, as in the manuscript eq. (7), is repeated here for convenience.

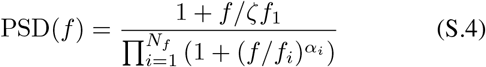

with *f*_*i*_ the *i*th cut-off frequency, *α*_*i*_ the attenuation coefficient and *N*_*f*_ = 2 or 3, for the vibrational and static interference analysis, respectively. *ζ* is randomly chosen between [0.3, 1]. Consequently, the PSD reaches a maximum between *ζf*_1_ and *f*_1_. This formulation was chosen based on the power-law (1*/f*^*α*^) power spectrum observed at many spatio-temporal scales inside the brain [41]–[43], [61]. The reported power-law coefficients in literature are typically fit for frequencies below 1 kHz. To obtain insides in the behavior at the higher frequencies (*>* 1 kHz). Dipole moments were calculated using morphologically accurate neuron models.

The used neuron models are multi-compartmental. Two hippocampal pyramidal models (morphologies: mpg141208_B_idA and mpg141209_A_idA) from Migliore et al. (2018) [45] were tested, and one L2/3 cortical pyramidal cell from Aberra et al. (2018) [44]. The models were obtained from ModelDB [62]. The accession numbers are 244688 and 241165, respectively. The transmembrane voltage V of a single compartment can be determined by:

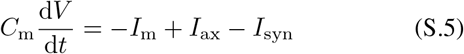

where *C*_m_ is the membrane capacitance, *I*_m_ the transmembrane current flowing out of the considered compartment, *I*_ax_ the axial current flowing into the considered compartment and *I*_syn_ an AMPA-like synaptic current. The latter is modeled as follows:

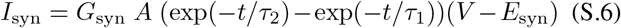

Synapses were randomly allocated to half of the apical dendritic compartments. A synaptic event was triggered every 2 ms. The rise time (*τ*_1_) equals 0.4 ms, the decay time (*τ*_2_) 1 ms and the weight *G*_syn_ 3 nS. *A* is a normalization factor in order to have a peak conductance of *G*_syn_. The simulations are performed with the NEURON simulation software [63] for 100 ms and with a fixed time step of 25 *μ*s. According to Murakimi et al. (2003) [46], the time dependent current dipole (**d**(*t*)) is calculated as follows:

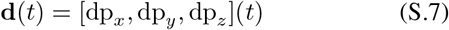

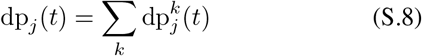

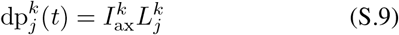

with 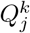 the current dipole and 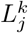 the length in direction *j* = *x, y, z* of compartment *k*.

Next, the PSD of the current dipole magnitude (|**d**| (*t*)) is determined. The frequency domain is divided into three regimes: low (*f ϵ* [0, 500 [*Hz*), medium (*f ϵ* [0.5, 5 [*kHz*) and high (*f ϵ* [5, 40 [*kHz*). A power-law is fit to each regime for each model via linear regression of the PSD at the log-log scale. The results and corresponding *α* coefficients are shown in figure S.1.

**Fig. S.1.**
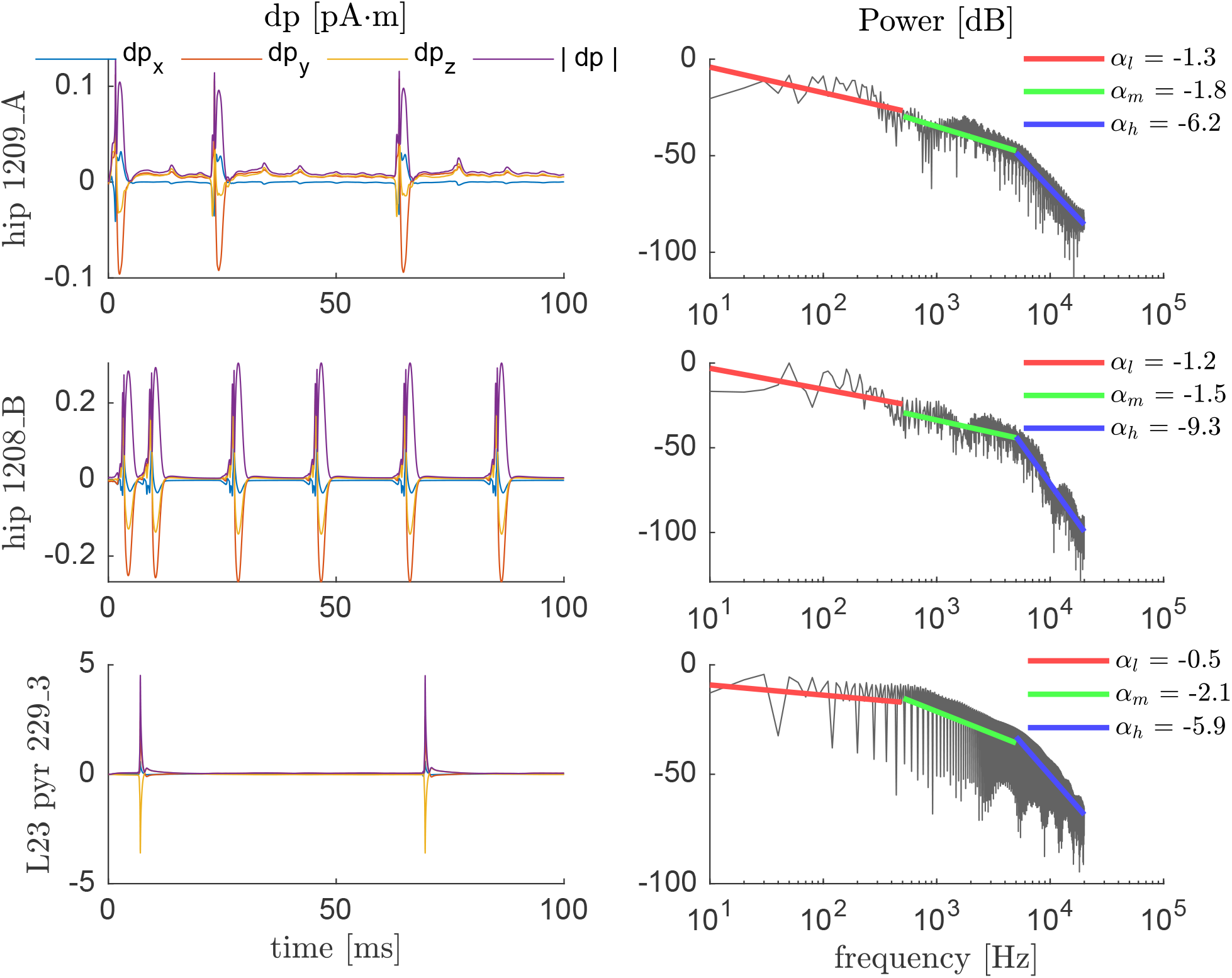
Dipole moment of tested models (y-label) in response to a pulsed synaptic input. (Left) signals in time domain, (right) corresponding power spectral density profile. The fitted power law coefficients for fixed frequency range are displayed in the legend. *α*_*l*_ : *fϵ*[0, 500] Hz, *α*_*m*_: *fϵ*[0.5, 5 [kHz and *α*_*h*_ :*fϵ*[5, 40 [kHz

## References

[1] P. Pinti, I. Tachtsidis, A. Hamilton, J. Hirsch, C. Aichelburg, S. Gilbert, and P. W. Burgess, “The present and future use of functional nearinfrared spectroscopy (fnirs) for cognitive neuroscience,” Annals of the New York Academy of Sciences, vol. 1464, pp. 5–29, 3 2020. [Online]. Available: https://onlinelibrary.wiley.com/doi/10.1111/nyas.13948

[2] S. Baillet, J. Mosher, and R. Leahy, “Electromagnetic brain mapping,” IEEE Signal Processing Magazine, vol. 18, pp. 14–30, 2001. [Online]. Available: http://ieeexplore.ieee.org/document/962275/

[3] B. He, A. Sohrabpour, E. Brown, and Z. Liu, “Electrophysiological source imaging: A noninvasive window to brain dynamics,” Annual Review of Biomedical Engineering, vol. 20, pp. 171–196, 6 2018. [Online]. Available: https://www.annualreviews.org/doi/10.1146/annurev-bioeng-062117-120853

[4] L. F. Nicolas-Alonso and J. Gomez-Gil, “Brain computer interfaces, a review,” Sensors, vol. 12, pp. 1211–1279, 1 2012. [Online]. Available: http://www.mdpi.com/1424-8220/12/2/1211

[5] S. Bollmann and M. Barth, “New acquisition techniques and their prospects for the achievable resolution of fmri,” Progress in Neurobiology, vol. 207, 12 2021. [Online]. Available: https://linkinghub.elsevier.com/retrieve/pii/S030100822030191X

[6] N. Logothetis, “Mr imaging in the non-human primate: studies of function and of dynamic connectivity,” Current Opinion in Neurobiology, vol. 13, pp. 630–642, 10 2003. [Online]. Available: https://linkinghub.elsevier.com/retrieve/pii/S0959438803001466

[7] R. P. Kennan, S. G. Horovitz, A. Maki, Y. Yamashita, H. Koizumi, and J. C. Gore, “Simultaneous recording of event-related auditory oddball response using transcranial near infrared optical topography and surface eeg,” NeuroImage, vol. 16, pp. 587–592, 7 2002. [Online]. Available: https://linkinghub.elsevier.com/retrieve/pii/S1053811902910608

[8] V. Quaresima and M. Ferrari, “Functional near-infrared spectroscopy (fnirs) for assessing cerebral cortex function during human behavior in natural/social situations: A concise review,” Organizational Research Methods, vol. 22, pp. 46–68, 1 2019. [Online]. Available: http://journals.sagepub.com/doi/10.1177/1094428116658959

[9] G. Buzsáki, C. A. Anastassiou, and C. Koch, “The origin of extracellular fields and currents — eeg, ecog, lfp and spikes,” Nature Reviews Neuroscience, vol. 13, pp. 407–420, 6 2012. [Online]. Available: http://www.nature.com/articles/nrn3241

[10] M. F. Hnazaee, E. Khachatryan, and M. M. V. Hulle, “Semantic features reveal different networks during word processing: An eeg source localization study,” Frontiers in Human Neuroscience, vol. 12, p. 503, 12 2018. [Online]. Available: https://www.frontiersin.org/article/10.3389/fnhum.2018.00503/full

[11] D. Sharon, M. S. Hämäläinen, R. B. Tootell, E. Halgren, and J. W. Belliveau, “The advantage of combining meg and eeg: Comparison to fmri in focally stimulated visual cortex,” NeuroImage, vol. 36, pp. 1225–1235, 7 2007. [Online]. Available: https://linkinghub.elsevier.com/retrieve/pii/S1053811907002133

[12] S. Baillet, “Magnetoencephalography for brain electrophysiology and imaging,” Nature Neuroscience, vol. 20, pp. 327–339, 3 2017. [Online]. Available: https://www.nature.com/articles/nn.4504

[13] S. Klamer, A. Elshahabi, H. Lerche, C. Braun, M. Erb, K. Scheffler, and N. K. Focke, “Differences between meg and high-density eeg source localizations using a distributed source model in comparison to fmri,” Brain Topography, vol. 28, pp. 87–94, 1 2015. [Online]. Available: http://link.springer.com/10.1007/s10548-014-0405-3

[14] A. K. Liu, A. M. Dale, and J. W. Belliveau, “Monte carlo simulation studies of eeg and meg localization accuracy,” Human Brain Mapping, vol. 16, pp. 47–62, 5 2002. [Online]. Available: https://onlinelibrary.wiley.com/doi/10.1002/hbm.10024

[15] A. Dubey and S. Ray, “Cortical electrocorticogram (ecog) is a local signal,” The Journal of Neuroscience, vol. 39, pp. 4299–4311, 5 2019. [Online]. Available: https://www.jneurosci.org/lookup/doi/10.1523/JNEUROSCI.2917-18.2019

[16] C. Todaro, L. Marzetti, P. A. V. Sosa, P. A. Valdés-Hernandez, and V. Pizzella, “Mapping brain activity with electrocorticography: Resolution properties and robustness of inverse solutions,” Brain Topography, vol. 32, pp. 583–598, 7 2019. [Online]. Available: http://link.springer.com/10.1007/s10548-018-0623-1

[17] S. Asadzadeh, T. Y. Rezaii, S. Beheshti, A. Delpak, and S. Meshgini, “A systematic review of eeg source localization techniques and their applications on diagnosis of brain abnormalities,” Journal of Neuroscience Methods, vol. 339, p. 108740, 6 2020. [Online]. Available: https://linkinghub.elsevier.com/retrieve/pii/S0165027020301631

[18] G. Birot, L. Spinelli, S. Vulliémoz, P. Mégevand, D. Brunet, M. Seeck, and C. M. Michel, “Head model and electrical source imaging: A study of 38 epileptic patients,” NeuroImage: Clinical, vol. 5, pp. 77–83, 2014. [Online]. Available: https://linkinghub.elsevier.com/retrieve/pii/S2213158214000813

[19] L. Koessler, C. Benar, L. Maillard, J.-M. Badier, J. P. Vignal, F. Bartolomei, P. Chauvel, and M. Gavaret, “Source localization of ictal epileptic activity investigated by high resolution eeg and validated by seeg,” NeuroImage, vol. 51, pp. 642–653, 6 2010. [Online]. Available: https://linkinghub.elsevier.com/retrieve/pii/S1053811910002545

[20] A. Sohrabpour, Y. Lu, P. Kankirawatana, J. Blount, H. Kim, and B. He, “Effect of eeg electrode number on epileptic source localization in pediatric patients,” Clinical Neurophysiology, vol. 126, pp. 472–480, 3 2015. [Online]. Available: https://linkinghub.elsevier.com/retrieve/pii/S1388245714003642

[21] H. Hallez, B. Vanrumste, R. Grech, J. Muscat, W. D. Clercq, A. Vergult, Y. D’Asseler, K. P. Camilleri, S. G. Fabri, S. V. Huffel, and I. Lemahieu, “Review on solving the forward problem in eeg source analysis,” Journal of NeuroEngineering and Rehabilitation, vol. 4, p. 46, 12 2007. [Online]. Available: https://jneuroengrehab.biomedcentral.com/articles/10.1186/1743-0003-4-46

[22] C. Rabut, M. Correia, V. Finel, S. Pezet, M. Pernot, T. Deffieux, and M. Tanter, “4d functional ultrasound imaging of whole-brain activity in rodents,” Nature Methods, vol. 16, pp. 994–997, 10 2019. [Online]. Available: http://www.nature.com/articles/s41592-019-0572-y

[23] B. He, “Focused ultrasound help realize high spatiotemporal brain imaging? - a concept on acousto-electrophysiological neuroimaging,” IEEE Transactions on Biomedical Engineering, vol. 63, pp. 2654–2656, 12 2016. [Online]. Available: https://ieeexplore.ieee.org/document/7676270/

[24] R. Olafsson, R. S. Witte, S.-W. Huang, and M. O’Donnell, “Ultrasound current source density imaging,” IEEE Transactions on Biomedical Engineering, vol. 55, pp. 1840–1848, 7 2008, afleiding UCSDI reconstruction. [Online]. Available: https://ieeexplore.ieee.org/document/4457869/

[25] R. Witte, R. Olafsson, S.-W. Huang, and M. O’Donnell, “Imaging current flow in lobster nerve cord using the acoustoelectric effect,” Applied Physics Letters, vol. 90, 4 2007. [Online]. Available: http://aip.scitation.org/doi/10.1063/1.2724901

[26] H. Zhang, M. Xu, M. Liu, X. Song, F. He, S. Chen, and D. Ming, “Biological current source imaging method based on acoustoelectric effect: A systematic review,” Frontiers in Neuroscience, vol. 16, 7 2022. [Online]. Available: https://www.frontiersin.org/articles/10.3389/fnins.2022.807376/full

[27] Z. H. Wang, R. Olafsson, P. Ingram, Q. Li, Y. Qin, and R. S. Witte, “Four-dimensional ultrasound current source density imaging of a dipole field,” Applied Physics Letters, vol. 99, 9 2011. [Online]. Available: http://aip.scitation.org/doi/10.1063/1.3632034

[28] R. Olafsson, R. S. Witte, C. Jia, S.-W. Huang, K. Kim, and M. O’donnell, “Cardiac activation mapping using ultrasound current source density imaging (ucsdi),” IEEE Transactions on Ultrasonics, Ferroelectrics, and Frequency Control, vol. 56, pp. 565–574, 3 2009. [Online]. Available: https://ieeexplore.ieee.org/document/4816064/

[29] Y. Qin, P. Ingram, A. Burton, and R. S. Witte, “4d acoustoelectric imaging of current sources in a human head phantom,” vol. 2016-November. IEEE, 9 2016, pp. 1–4. [Online]. Available: http://ieeexplore.ieee.org/document/7728868/

[30] A. Barragan, C. Preston, A. Alvarez, T. Bera, Y. Qin, M. Weinand, W. Kasoff, and R. S. Witte, “Acoustoelectric imaging of deep dipoles in a human head phantom for guiding treatment of epilepsy,” Journal of Neural Engineering, vol. 17, 10 2020. [Online]. Available: https://iopscience.iop.org/article/10.1088/1741-2552/abb63a

[31] R. M. Arthur and D. B. Geselowitz, “Effect of inhomogeneities on the apparent location and magnitude of a cardiac current dipole source,” IEEE Transactions on Biomedical Engineering, vol. BME-17, pp. 141–146, 4 1970. [Online]. Available: http://ieeexplore.ieee.org/document/4502713/

[32] Y. Salu, L. Cohen, D. Rose, S. Sxato, C. Kufta, and M. Hallett, “An improved method for localizing electric brain dipoles,” IEEE Transactions on Biomedical Engineering, vol. 37, pp. 699–705, 7 1990. [Online]. Available: http://ieeexplore.ieee.org/document/55680/

[33] P. Hasgall, F. D. Gennaro, C. Baumgartner, E. Neufeld, B. Lloyd, M. Gosselin, D. Payne, A. Klingenböck, and N. Kuster, “It’is database for thermal and electromagnetic parameters of biological tissues,” vol. 4.0, 5 2018.

[34] S. Murakami and Y. Okada, “Invariance in current dipole moment density across brain structures and species: Physiological constraint for neuroimaging,” NeuroImage, vol. 111, pp. 49–58, 5 2015.

[35] D. P. Buxhoeveden and M. F. Casanova, “The minicolumn hypothesis in neuroscience,” Brain, vol. 125, pp. 935–951, 5 2002. [Online]. Available: https://academic.oup.com/brain/article-lookup/doi/10.1093/brain/awf110

[36] C. for Devices and R. Health, “Food and drug administration, u.s. department of health and human services” 6.

[37] H. T. O’Neil, “Theory of focusing radiators,” The Journal of the Acoustical Society of America, vol. 21, pp. 516–526, 9 1949. [Online]. Available: http://asa.scitation.org/doi/10.1121/1.1906542

[38] J. K. Mueller, L. Ai, P. Bansal, and W. Legon, “Computational exploration of wave propagation and heating from transcranial focused ultrasound for neuromodulation,” Journal of Neural Engineering, vol. 13, 10 2016. [Online]. Available: https://iopscience.iop.org/article/10.1088/1741-2560/13/5/056002

[39] D. N. Stephens, D. E. Kruse, S. Qin, and K. W. Ferrara, “Design aspects of focal beams from high-intensity arrays,” IEEE Transactions on Ultrasonics, Ferroelectrics and Frequency Control, vol. 58, pp. 1590–1602, 8 2011. [Online]. Available: http://ieeexplore.ieee.org/document/5995216/

[40] S. Murakami and Y. Okada, “Contributions of principal neocortical neurons to magnetoencephalography and electroencephalography signals,” The Journal of Physiology, vol. 575, pp. 925–936, 9 2006, info on dipole moment strength. [Online]. Available: https://onlinelibrary.wiley.com/doi/10.1113/jphysiol.2006.105379

[41] B. J. He, “Scale-free brain activity: past, present, and future,” Trends in Cognitive Sciences, vol. 18, pp. 480–487, 9 2014. [Online]. Available: https://linkinghub.elsevier.com/retrieve/pii/S1364661314000850

[42] K. H. Pettersen, H. Lindén, T. Tetzlaff, and G. T. Einevoll, “Power laws from linear neuronal cable theory: Power spectral densities of the soma potential, soma membrane current and single-neuron contribution to the eeg,” PLoS Computational Biology, vol. 10, 11 2014. [Online]. Available: https://dx.plos.org/10.1371/journal.pcbi.1003928

[43] S. X. Moffett, S. M. O’Malley, S. Man, D. Hong, and J. V. Martin, “Dynamics of high frequency brain activity,” Scientific Reports, vol. 7, 12 2017. [Online]. Available: http://www.nature.com/articles/s41598-017-15966-6

[44] A. S. Aberra, A. V. Peterchev, and W. M. Grill, “Biophysically realistic neuron models for simulation of cortical stimulation,” Journal of Neural Engineering, vol. 15, 12 2018. [Online]. Available: https://iopscience.iop.org/article/10.1088/1741-2552/aadbb1

[45] R. Migliore, C. A. Lupascu, L. L. Bologna, A. Romani, J.-D. Courcol, S. Antonel, W. A. H. V. Geit, A. M. Thomson, A. Mercer, S. Lange, J. Falck, C. A. Rössert, Y. Shi, O. Hagens, M. Pezzoli, T. F. Freund, S. Kali, E. B. Muller, F. Schürmann, H. Markram, and M. Migliore, “The physiological variability of channel density in hippocampal ca1 pyramidal cells and interneurons explored using a unified data-driven modeling workflow,” PLOS Computational Biology, vol. 14, 9 2018. [Online]. Available: https://dx.plos.org/10.1371/journal.pcbi.1006423

[46] S. Murakami, A. Hirose, and Y. C. Okada, “Contribution of ionic currents to magnetoencephalography (meg) and electroencephalography (eeg) signals generated by guinea-pig ca3 slices,” The Journal of Physiology, vol. 553, pp. 975–985, 12 2003.

[47] Q. Li, R. Olafsson, P. Ingram, Z. Wang, and R. Witte, “Measuring the acoustoelectric interaction constant using ultrasound current source density imaging,” Physics in Medicine and Biology, vol. 57, pp. 5929–5941, 10 2012. [Online]. Available: https://iopscience.iop.org/article/10.1088/0031-9155/57/19/5929

[48] J. Zimmermann and U. van Rienen, “Ambiguity in the interpretation of the low-frequency dielectric properties of biological tissues,” Bioelectrochemistry, vol. 140, 8 2021. [Online]. Available: https://linkinghub.elsevier.com/retrieve/pii/S1567539421000360

[49] E. Mehić, J. M. Xu, C. J. Caler, N. K. Coulson, C. T. Moritz, and P. D. Mourad, “Increased anatomical specificity of neuromodulation via modulated focused ultrasound,” PLoS ONE, vol. 9, 2 2014. [Online]. Available: https://dx.plos.org/10.1371/journal.pone.0086939

[50] W. Legon, T. F. Sato, A. Opitz, J. Mueller, A. Barbour, A. Williams, and W. J. Tyler, “Transcranial focused ultrasound modulates the activity of primary somatosensory cortex in humans,” Nature Neuroscience, vol. 17, pp. 322–329, 2 2014.

[51] D. P. Darrow, “Focused ultrasound for neuromodulation,” Neurotherapeutics, vol. 16, pp. 88–99, 1 2019.

[52] T. Lemaire, E. Neufeld, N. Kuster, and S. Micera, “Understanding ultrasound neuromodulation using a computationally efficient and interpretable model of intramembrane cavitation,” Journal of Neural Engineering, vol. 16, p. 046007, 7 2019.

[53] T. Tarnaud, W. Joseph, R. Schoeters, L. Martens, and E. Tanghe, “Seconic: Towards multi-compartmental models for ultrasonic brain stimulation by intramembrane cavitation,” Journal of Neural Engineering, vol. 17, p. 056010, 10 2020. [Online]. Available: https://iopscience.iop.org/article/10.1088/1741-2552/abb73d

[54] T. Tarnaud, W. Joseph, R. Schoeters, L. Martens, and E. Tanghe, “Improved alpha-beta power reduction via combined electrical and ultrasonic stimulation in a parkinsonian cortex-basal ganglia-thalamus computational model,” Journal of Neural Engineering, vol. 18, 12 2021. [Online]. Available: https://iopscience.iop.org/article/10.1088/1741-2552/ac3f6d

[55] T. Kim, C. Park, P. Y. Chhatbar, J. Feld, B. M. Grory, C. S. Nam, P. Wang, M. Chen, X. Jiang, and W. Feng, “Effect of low intensity transcranial ultrasound stimulation on neuromodulation in animals and humans: An updated systematic review,” Frontiers in Neuroscience, vol. 15, 4 2021. [Online]. Available: https://www.frontiersin.org/articles/10.3389/fnins.2021.620863/full

[56] M. Plaksin, E. Kimmel, and S. Shoham, “Cell-type-selective effects of intramembrane cavitation as a unifying theoretical framework for ultrasonic neuromodulation,” eneuro, vol. 3, 5 2016. [Online]. Available: https://www.eneuro.org/lookup/doi/10.1523/ENEURO.0136-15.2016

[57] F. Darvas, E. Mehić, C. J. Caler, J. G. Ojemann, and P. D. Mourad, “Toward deep brain monitoring with superficial eeg sensors plus neuromodulatory focused ultrasound,” Ultrasound in Medicine and Biology, vol. 42, pp. 1834–1847, 8 2016. [Online]. Available: https://linkinghub.elsevier.com/retrieve/pii/S0301562916001137

[58] T. Tarnaud, W. Joseph, L. Martens, and E. Tanghe, “Computational modeling of ultrasonic subthalamic nucleus stimulation,” IEEE Transactions on Biomedical Engineering, vol. 66, pp. 1155–1164, 4 2019. [Online]. Available: https://ieeexplore.ieee.org/document/8456629/

[59] T. Sato, M. G. Shapiro, and D. Y. Tsao, “Ultrasonic neuromodulation causes widespread cortical activation via an indirect auditory mechanism,” Neuron, vol. 98, pp. 1031–1041.e5, 6 2018. [Online]. Available: https://linkinghub.elsevier.com/retrieve/pii/S0896627318303817

[60] H. Guo, M. Hamilton, S. J. Offutt, C. D. Gloeckner, T. Li, Y. Kim, W. Legon, J. K. Alford, and H. H. Lim, “Ultrasound produces extensive brain activation via a cochlear pathway,” Neuron, vol. 98, pp. 1020–1030.e4, 6 2018. [Online]. Available: https://linkinghub.elsevier.com/retrieve/pii/S0896627318303714

[61] G. Buzsáki and K. Mizuseki, “The log-dynamic brain: how skewed distributions affect network operations,” Nature Reviews Neuroscience, vol. 15, pp. 264–278, 4 2014. [Online]. Available: http://www.nature.com/articles/nrn3687

[62] R. A. McDougal, T. M. Morse, T. Carnevale, L. Marenco, R. Wang, M. Migliore, P. L. Miller, G. M. Shepherd, and M. L. Hines, “Twenty years of modeldb and beyond: building essential modeling tools for the future of neuroscience,” Journal of Computational Neuroscience, vol. 42, pp. 1–10, 2 2017.

[63] N. T. Carnevale and M. L. Hines, The NEURON book. Cambridge University Press, 2006.

